# Cerebellar learning using perturbations

**DOI:** 10.1101/053785

**Authors:** Guy Bouvier, Johnatan Aljadeff, Claudia Clopath, Célian Bimbard, Jonas Ranft, Antonin Blot, Jean-Pierre Nadal, Nicolas Brunel, Vincent Hakim, Boris Barbour

## Abstract

The cerebellum aids the learning and execution of fast coordinated movements, with acquired information being stored by plasticity of parallel fibre—Purkinje cell synapses. According to the current consensus, erroneously active parallel fibre synapses are depressed by complex spikes arising when climbing fibres signal movement errors. However, this theory cannot solve the *credit assignment problem* of using the limited information from a global movement evaluation to optimise behaviour by guiding the plasticity in numerous neurones. We identify the possible implementation of an algorithm solving this problem, whereby spontaneous complex spikes perturb ongoing movements, create an eligibility trace for plasticity and signal resulting error changes to guide plasticity. These error changes are extracted by adaptively cancelling the average error. This framework, *stochastic gradient descent with estimated global errors*, generates specific predictions for synaptic plasticity rules that contradict the current consensus. However, in vitro plasticity experiments under physiological conditions verified our predictions, highlighting the sensitivity of plasticity studies to unphysiological conditions. Using numerical and analytical approaches we demonstrate the convergence and estimate the capacity of learning in our implementation. Finally, a similar mechanism may operate during optimisation of action sequences by the basal ganglia, where dopamine could both initiate movements and signal rewards, analogously to the dual perturbation and correction role of the climbing fibre outlined here.

## 1 Introduction

A central contribution of the cerebellum to motor control is thought to be the learning and automatic execution of fast, coordinated movements. Anatomically, the cerebellum consists of a convoluted, lobular cortex surrounding the cerebellar nuclei (Fig. 1A). The main input to the cerebellum is the heterogeneous mossy fibres, which convey multiple modalities of sensory, contextual and motor information. They excite both the cerebellar nuclei and the cerebellar cortex; in the cortex they synapse with the very abundant granule cells, whose axons, the parallel fibres, excite Purkinje cells. Purkinje cells constitute the sole output of the cerebellar cortex and project an inhibitory connection to the nuclei, which therefore combine a direct and a transformed mossy fibre input with opposite signs. The largest cell type in the nuclei, the projection neurones, send excitatory axons to several motor effector systems, notably the motor cortex via the thalamus. Another nuclear cell type, the nucleo-olivary neurones, inhibit the inferior olive. The cerebellum receives a second external input: climbing fibres from the inferior olive, which form an extensive, ramified connection with the proximal dendrites of the Purkinje cell. Each Purkinje cell receives a single climbing fibre. A more modular diagram of the olivo-cerebellar connectivity relevant to this paper is shown in Fig. 1B; numerous cell types and connections have been omitted for simplicity.

**Figure 1:**
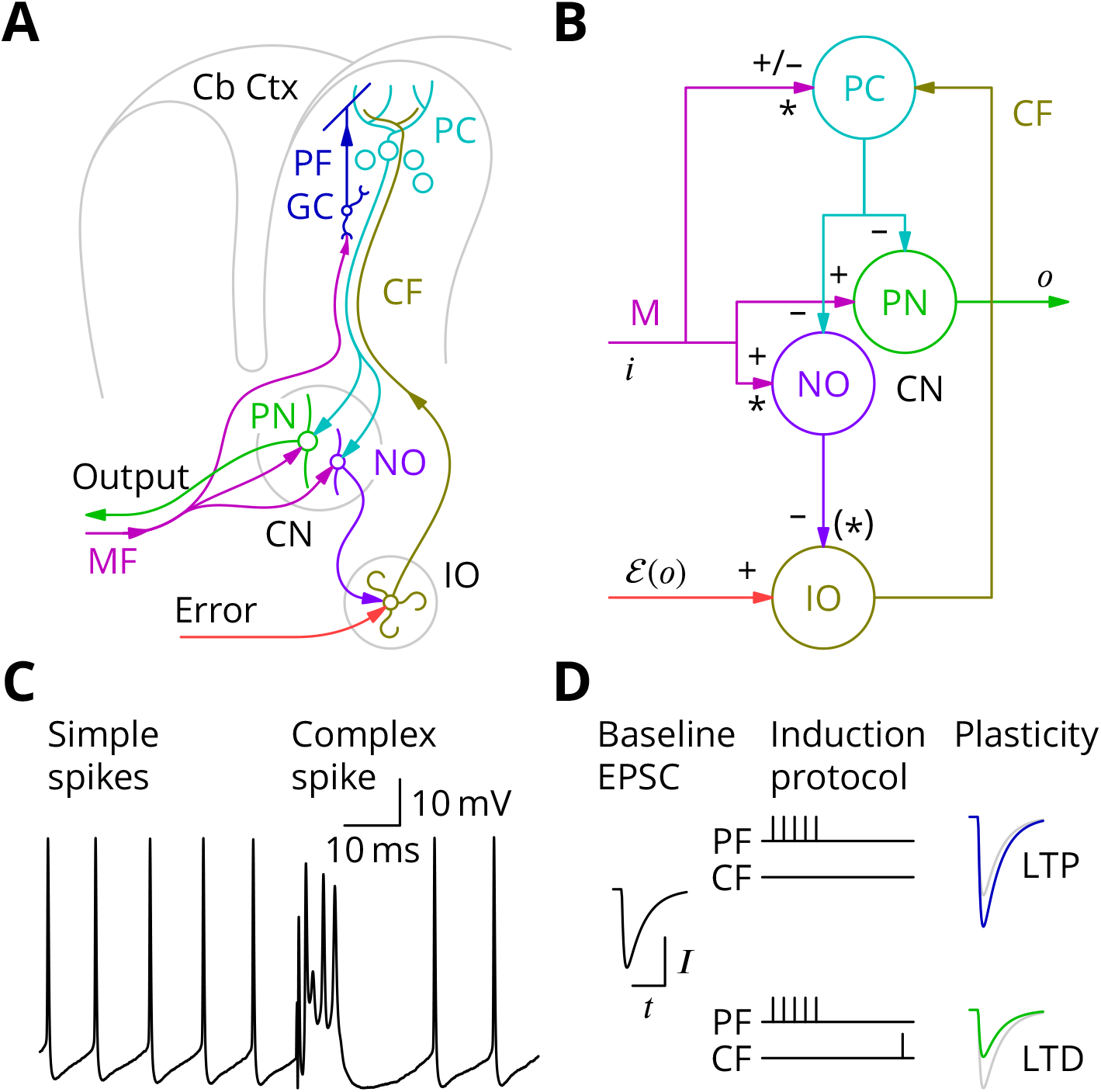
The cerebellar circuitry and properties of Purkinje cells. **A**. Simplified circuit diagram. MF, mossy fibres; CN, (deep) cerebellar nuclei; GC, granule cells; Cb Ctx, cerebellar cortex; PF, parallel fibres; PC, Purkinje cells; PN, projection neurones; NO, nucleo-olivary neurones; IO, inferior olive; CF, Climbing fibres. **B**. Modular diagram. The signs of the synapses are indicated. The granule cell and indirect inhibitory inputs they recruit have been subsumed into a bidirectional mossy fibre–Purkinje cell input, M. Potentially plastic inputs of interest here are denoted with an asterisk. *i*, input; *o*, output; 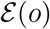, error (which is a function of the output). **C**. Typical Purkinje cell electrical activity from an intracellular patch-clamp recording. Purkinje cells fire two types of action potential: simple spikes and, in response to climbing fibre input, complex spikes. **D**. According to the consensus plasticity rule, a complex spike will depress parallel fibre synapses active about 100 ms earlier. The diagram depicts idealised excitatory postsynaptic currents (EPSCs) before and after typical induction protocols inducing long-term potentiation (LTP) or depression (LTD). Grey, control EPSC; blue, green, post-induction EPSCs.

Purkinje cells discharge two distinct types of action potential (Fig. 1C). They nearly continuously emit *simple spikes*—standard, if brief, action potentials—at frequencies that average 50 Hz. This frequency is modulated both positively and negatively by the intensity of inputs from the mossy fibre–granule cell pathway (which can also recruit interneurones that inhibit Purkinje cells; Eccles et al., 1967). Such modulations of Purkinje cell firing are thought to underlie their contributions to motor control. In addition, when the climbing fibre is active, an event that occurs continuously but in a somewhat irregular pattern with a mean frequency of around 1 Hz, the Purkinje cell emits a completely characteristic *complex spike* under the influence of the intense excitation from the climbing fibre (Fig. 1C).

The history of research into cerebellar learning is dominated by the theory due to Marr (1969) and Albus (1971). They suggested that the climbing fibre acts as a ‘teacher’ to guide plasticity of parallel fibre–Purkinje cell synapses. It was several years, however, before experimental support for this hypothesis was obtained (Ito et al., 1982; Ito and Kano, 1982), while the notion that the climbing fibre signalled errors had emerged by that time (Ito, 1972, 1984). Error modalities thought to be represented by climbing fibres include: pain, unexpected touch, imbalance, and retinal slip. According to the modern understanding of this theory, by signalling such movement errors, climbing fibres induce long-term depression (LTD) of parallel fibre synapses that were active at the same time (Ito et al., 1982; Ito and Kano, 1982; Sakurai, 1987; Crepel and Jaillard, 1991) or, more precisely, shortly before (Wang et al., 2000; Sarkisov and Wang, 2008; Safo and Regehr, 2008). A compensating long-term potentiation (LTP) is necessary to prevent synaptic saturation (Lev-Ram et al., 2002, 2003; Coesmans et al., 2004) and its induction is reported to follow high-frequency parallel fibre activity in the absence of complex spikes (Jörntell and Ekerot, 2002; Bouvier et al., 2016). Plasticity of parallel fibre synaptic currents according to these plasticity rules is diagrammed in Fig. 1D.

The Marr-Albus-Ito theory represents a form of supervised learning—in which an external teacher provides the correct output. The theory is incomplete, however, as it does not describe how a global evaluation of movement error can be processed to provide cell-specific instructions for plasticity to large numbers of cells. In fact, the inverse problem of deducing the precise synaptic changes required to correct an error within a large circuit controlling complex, arbitrary tasks is for all practical purposes intractable, a situation for which Minsky (1961) coined the term the *credit assignment problem* (where *credit* implies maximisation of a reward rather than the mathematically equivalent minimisation of an error). But, for the following reasons, it is likely that the cerebellum *is* able to solve this problem. The ability to learn arbitrary sensory-motor associations would offer great advantage to an organism. Conversely, application of an inadequate learning algorithm would have catastrophic behavioural consequences, with the ‘learning’ potentially reinforcing the error-producing action. However, such wrong-headed learning seems never to be observed, strongly suggesting that the cerebellum does implement an algorithm able to avoid this problem.

A suitable algorithm for solving the general cerebellar learning problem would be *stochastic gradient descent* (Minsky, 1961), according to which the objective function is explored by random variations in the network that alter behaviour, with plasticity then retaining those variations that improve the behaviour, as signalled by a decreased error or increased reward. Several possible mechanisms of varying biological plausibility have been proposed. In particular, perturbations caused by synaptic release (Minsky, 1954; Seung, 2003) or external inputs (Doya and Sejnowski, 1988) have been suggested, while extraction of changes in the objective function can, for instance, be performed implicitly using a scheme proposed by Williams (1992). Although the theoretical framework for gradient descent is well established, the goal of identifying in the brain a network and cellular implementation of such an algorithm has proved elusive.

The learning behaviour with the best established resemblance to stochastic gradient ascent is the acquisition of song in male songbirds. The juvenile song is refined by a trial and error process to approach a template memorised from a tutor during a critical period (Konishi, 1965; Mooney, 2009). The analogy with stochastic gradient ascent was made by Doya and Sejnowski (1988) and was then further developed experimentally (Olveczky et al., 2005) and theoretically (Fiete et al., 2007). However, despite these very suggestive behavioural correlates, relatively little progress has been made in verifying model predictions for plasticity or identifying the structures responsible for storing the template and evaluating its match with the song.

A gradient descent mechanism for the cerebellum has been proposed by the group of Dean et al. (2002), who term their algorithm *decorrelation*. Simply stated (Dean and Porrill, 2014), if both parallel fibre and climbing fibre inputs to a Purkinje cell are assumed to vary about their respective mean values, their correlation or anticorrelation indicates the local gradient of the error function and thus the sign of the required plasticity. At the optimum, which is a minimum of the error function, there should be no correlation between variations of the climbing fibre rate and those of the parallel fibre input. Hence the name: the algorithm aims to decorrelate parallel and climbing fibre variations^1^. An appropriate plasticity rule is a modified covariance rule (Sejnowski, 1977, who moreover suggested in abridged form a similar application to cerebellar learning). Although decorrelation provides a suitable framework, its proponents are still in the process of developing a cellular implementation (Menzies et al., 2010).

In summary, current theory contains a very simple plasticity rule but probably requires unrealistically sophisticated evaluation of motor errors (below we shall simulate tasks that that the current theory would be incapable of optimising). We sought to identify in the cerebellar circuitry the implementation an algorithm transferring some of the algorithmic complexity from the evaluation to the plasticity rule. Below we propose a cellular implementation of stochastic gradient descent in the cerebellum that can achieve this goal. The central feature of our implementation is that the climbing fibre plays a dual role. In addition to its traditional function of conveying error information, we propose that the inferior olive perturbs movements by producing spontaneous complex spikes in Purkinje cells (Harris, 1998) and simultaneously creates an eligibility trace specific to the perturbed cells. That trace is then exploited using error signalling by the climbing fibre.

### 1.1 Article outline

We outline the organisation of this article. Readers should be aware that, although it contains an important experimental verification of a central prediction of our hypothesis, a large part of the article is given over to conceptual analysis, modelling and mathematical investigation of the algorithm we propose and its potential consequences for our understanding of cerebellar function.

We begin with an analysis of the requirements (§3) for cerebellar learning and of the perceived deficiencies of the current theory. We then develop an alternative algorithm based upon stochastic gradient descent, leading to the proposal that spontaneous complex spikes represent perturbations in a trial-and-error process (§4). We describe how this generates predictions for plasticity rules in Purkinje cells that contradict the current literature. There follows an experimental verification of these predictions (§5). We then suggest an implementation for an additional plasticity process required by the algorithm to extract the change of error (§6). A network simulation demonstrates the functionality of the algorithmic implementation (§7), while a more technical section (§8) establishes the convergence criteria and explores numerically the storage capacity of the algorithm. Finally, in the Discussion, we place our implementation in the context of the cerebellar literature and identify possible analogous mechanisms in other brain regions and preparations (§9).

## 2 Methods

### 2.1 Electrophysiology

Animal experimentation methods were authorised by the ‘Charles Darwin N°5’ ethics committee. Adult female C57Bl/6 mice (2–5 months old) were anæsthetised with isoflurane (Nicholas Piramal Ltd, India) and killed by decapitation. The cerebellum was rapidly dissected into cold solution containing (in mM): 230 sucrose, 26 NaHCO_3_, 3 KCl, 0.8 CaCl_2_, 8 MgCl_2_, 1.25 NaH_2_PO_4_, 25 d-glucose supplemented with 50 *μ*m d-APV to protect the tissue during slicing. 300 *μ*m sagittal slices were cut in this solution using a Campden Instruments 7000smz and stored at 32 °C in a standard extracellular saline solution containing: 125 NaCl, 2.5 KCl, 1.5 CaCl_2_, 1.8 MgCl_2_, 1.25 NaH_2_PO_4_, 26 NaHCO_3_ and 25 glucose, bubbled with 95% O_2_ and 5% CO_2_ (pH 7.4). Slices were visualised using an upright microscope with a 40 X, 0.8 NA water-immersion objective and infrared optics (illumination filter 750 ± 50 nm). The recording chamber was continuously perfused at a rate of 4–6 ml min^−1^ with a solution containing (mM): 125 NaCl, 2.5 KCl, 1.5 CaCl_2_, 1.8 MgCl_2_, 1.25 NaH_2_PO_4_, 26 NaHCO_3_, 25 d-glucose and 10 tricine, a Zn^2+^ buffer (Paoletti et al., 1997), bubbled with 95 % O_2_ and 5 % CO_2_ (pH 7.4). Patch pipettes had resistances in the range 2–4 MΩ with the internal solutions given below. Unless otherwise stated, cells were voltage clamped at −70 mV in the whole-cell configuration. Voltages are reported without correction for the junction potential, which was about 10 mV (so true membrane potentials were more negative than we report). Series resistances were 4–10 MΩ and compensated with settings of ~ 90 % in a Multiclamp 700B amplifier (Molecular Devices). Whole-cell recordings were filtered at 2kHz and digitised at 10 kHz. Experiments were performed at 32–34 °C. The internal solution contained (in mM): 128 K-gluconate, 10 HEPES, 4 KCl, 2.5 K_2_HPO_4_, 3.5 Mg-ATP, 0.4 Na_2_-GTP, 0.5 l-(–)-malic acid, 0.008 oxaloacetic acid, 0.18 *α*-ketoglutaric acid, 0.2 pyridoxal 5’-phosphate, 5 l-alanine, 0.15 pyruvic acid, 15 l-glutamine, 4 l-asparagine, 1 reduced l-glutathione, 0.5 NAD^+^, 5 phosphocreatine, 1.9 CaCl_2_, 1.5 MgCl_2_, K_3.8_EGTA. Free [Ca^2+^] was calculated with Maxchelator (C. Patton, Stanford) to be 120 nM. Chemicals were purchased from Sigma-Aldrich, d-APV from Tocris.

Recordings were made in the vermis of lobules three to eight of the cerebellar cortex. Granule cell EPSCs were elicited with stimulation in the granule cell layer with a glass pipette of tip diameter 8–12 *μ*m filled with HEPES-buffered saline. Climbing fibre electrodes had ~ 2 *μ*m diameter and were also positioned in the granule cell layer. Images were taken every 5 min; experiments showing significant slice movement (> 20*μ*m) were discarded. Stimulation intensity was fixed at the beginning of the experiment (1–15 V; 50–200 *μ*s) and maintained unchanged during the experiment. Test stimulation was applied at 0.1 Hz for the granule cell input and climbing fibres were stimulated at 0.5 Hz during the entire recording, mimicking tonic activity of the inferior olive in vivo (albeit at a slightly lower frequency). Test granule cell stimulation consisted of two pulses separated by 50 ms, allowing the quantification of paired-pulse facilitation. Test climbing fibre stimulation consisted of two pulses separated by 2.5 ms (400 Hz). The plasticity induction protocol involved stimulating granule cells with 5 pulses at 200 Hz every two seconds for 10 min with various relative timings of climbing fibre stimuli: a pair of climbing fibre stimuli at 400 Hz, 11–15 ms or ~ 500 ms after the start of the granule cell burst and/or four climbing fibre stimuli at 400 Hz, 100–115 ms after the beginning of the granule cell burst (timing diagrams will be shown in the Results). During induction, Purkinje cells were recorded in current-clamp mode with zero holding current.

### 2.2 Analysis

No formal prior power calculation to determine the sample sizes was performed; experiments were undertaken with the since verified expectation that interpretable results would require larger numbers of recordings than usual in plasticity studies (*n* < 10), because of the low signal-to-noise ratio of responses to stimulation in the granule cell layer. Recordings were analysed from 75 cells in 55 animals. Slices were systematically changed after each recording. Animals supplied 1–3 cells to the analysis, usually with rotating induction protocols. There was no blinding. Extending our modelling results below using a mixed-effect model (call lmer in the lme4 R package) to include animals as a random effect did not indicate that it was necessary to take into account any between-animal variance, as this was reported to be zero. On this basis, we consider each cell recorded to be the biological replicate in what follows.

Inspection of acquired climbing fibre responses revealed some failures of secondary stimuli in a fraction of cells, presumably because the second and subsequent stimuli at short intervals fell within the relative refractory period. As a complex spike was always produced these cells have been included in our analysis, but where individual data are displayed, we identify those cells in which failures of secondary climbing fibre stimuli were observed before the end of the induction period.

Analysis made use of Python scripts developed in house by Antonin Blot. Analysis of EPSC amplitudes began by averaging all of the EPSCs acquired to give a smooth time course. The time of the peak of this ‘global’ average response was determined. Subsequent measurement of amplitudes of other averages or of individual responses was performed by averaging the current over 0.5 ms centred on the time of the peak of the global average. The baseline calculated over 5 ms shortly before the stimulus artefact was subtracted to obtain the EPSC amplitude. Similar independent analyses were performed for both EPSCs of the paired-pulse stimulation.

Individual EPSCs were excluded from further analysis if the baseline current in the sweep exceeded −1 nA at −70 mV. Similarly, the analysis automatically excluded EPSCs in sweeps in which the granule cell stimulation elicited an action potential in the Purkinje cell (possibly through antidromic stimulation of its axon or through capacitive coupling to the electrode). However, during induction, in current clamp, such spikes were accepted. For displaying time series, granule cell responses were averaged in bins of 2 minutes.

The effects of series resistance changes in the Purkinje cell were estimated by monitoring the transient current flowing in response to a voltage step. The amplitude of the current 2 ms after the beginning of the capacity transient was measured. We shall call this the ‘dendritic access’. Modelling of voltage-clamp EPSC recordings in a two-compartment model of a Purkinje cell (Llano et al., 1991) suggests that this measure is approximately proportional to EPSC amplitude as the series resistance changes over a reasonable range (not shown). It therefore offers a better estimate of the effect on the EPSC amplitude of series resistance changes than would the value of the series resistance (or conductance), which is far from proportional to EPSC amplitude. Intuitively, this can be seen to arise because the EPSC is filtered by the dendritic compartment and the measure relates to the dendritic component of the capacitive transient, whereas the series resistance relates to the somatic compartment. We therefore calculated *R*_res_, the ratio of the dendritic access after induction (when plasticity was assessed) to the value before induction, in order to predict the changes the EPSC amplitude arising from changes of series resistance.

Because we elicited EPSCs using constant-voltage stimulation, variations of the resistance of the tip of the stimulating electrode (for instance if cells are drawn into it) could alter the stimulating current flow in the tissue. We monitored this by measuring the amplitude of the stimulus artefact. Specifically, we calculated *R*_stim_, the after/before ratio of the stimulation artefact amplitude.

We then used a robust linear model to examine the extent to which changes of series resistance or apparent stimulation strength could confound our measurements of plasticity, which we represented as the after/before ratio of EPSC amplitudes *R*_EPSC_; the model (in R syntax) was:

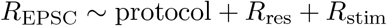

This showed that series resistance changes, represented by *R*_res_, had a significant influence (*t*-value 2.28, 69 degrees of freedom) with a slope close to the predicted unity (1.13). In contrast, changes of the stimulus artefact had no predictive value (slope −0.0008, *t*-value −0.003).

We did not wish to rely on the parametric significance tests of the linear model for comparing the plasticity protocols (although all of comparisons we report below as significant were also significant in the model). Instead, we equalised the dendritic filtering and stimulation changes between groups by eliminating those cells in which *R*_res_ or *R*_stim_ differed by more than 20 % from the mean values for all cells (0.94 ± 0.10 and 1.01 ± 0.19, respectively; mean ± sd, *n* = 75). After this operation, which eliminated 17 cells out of 75 leaving 58 (from 47 animals), the mean ratios varied by only a few percent between groups (ranges 5 % and 2 % for *R*_res_ and *R*_stim_, respectively) and would be expected to have only a minimal residual influence. Normalising the *R*_EPSC_s of the trimmed groups by *R*_res_ did not alter the conclusions presented below. After this trimming, the differences of *R*_EPSC_ between induction protocols were evaluated statistically using two-tailed nonparametric tests implemented by the wilcox.test command in R (R Core Team, 2013). Note that the remaining changes of *R*_res_ imply that all EPSC amplitudes after induction were underestimated by about 6 % relative to those at the beginning of the recording.

95 % confidence limits were calculated using R bootstrap functions (“BCa” method). For confidence limits of differences between means, stratified resampling was used.

### 2.3 Simulation methods

The model simulated in §7 was designed as follows (see also diagram in Fig. 2). A total of *S* × *L* Purkinje cells were placed on a rectangular grid of extent *S* in the sagittal plane and width *L* in the lateral direction. The activity of each Purkinje cell during a ‘movement’ was characterised by its firing rate

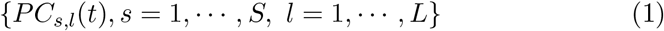

in *T* time bins {*t* = 1, ⋯, *T*}. Purkinje cells contacted *L* projection neurones (*PN*) in the cerebellar nuclei, which also contained *L* nucleo-olivary neurones (*NO*). The activities of both types of nuclear cells were also characterised by their firing rates in the different time bins, {*PN_l_*(*t*), *l* = 1, ⋯, *L*} and {*NO_l_*(*t*), *l* = 1, ⋯, *L*}. Mossy fibres, granule cells (parallel fibres) and molecular layer interneurones were subsumed into a single cell type (*M*) with *N* cells restricted to each row of *L* Purkinje cells, with a total of *N* × *S* mossy fibres. Mossy fibre activity was represented in a binary manner, *M_i,s_*(*t*) = 0 or 1, with {*i* = 1, ⋯, *N*} and {*s* = 1, ⋯, *S*}.

**Figure 2:**
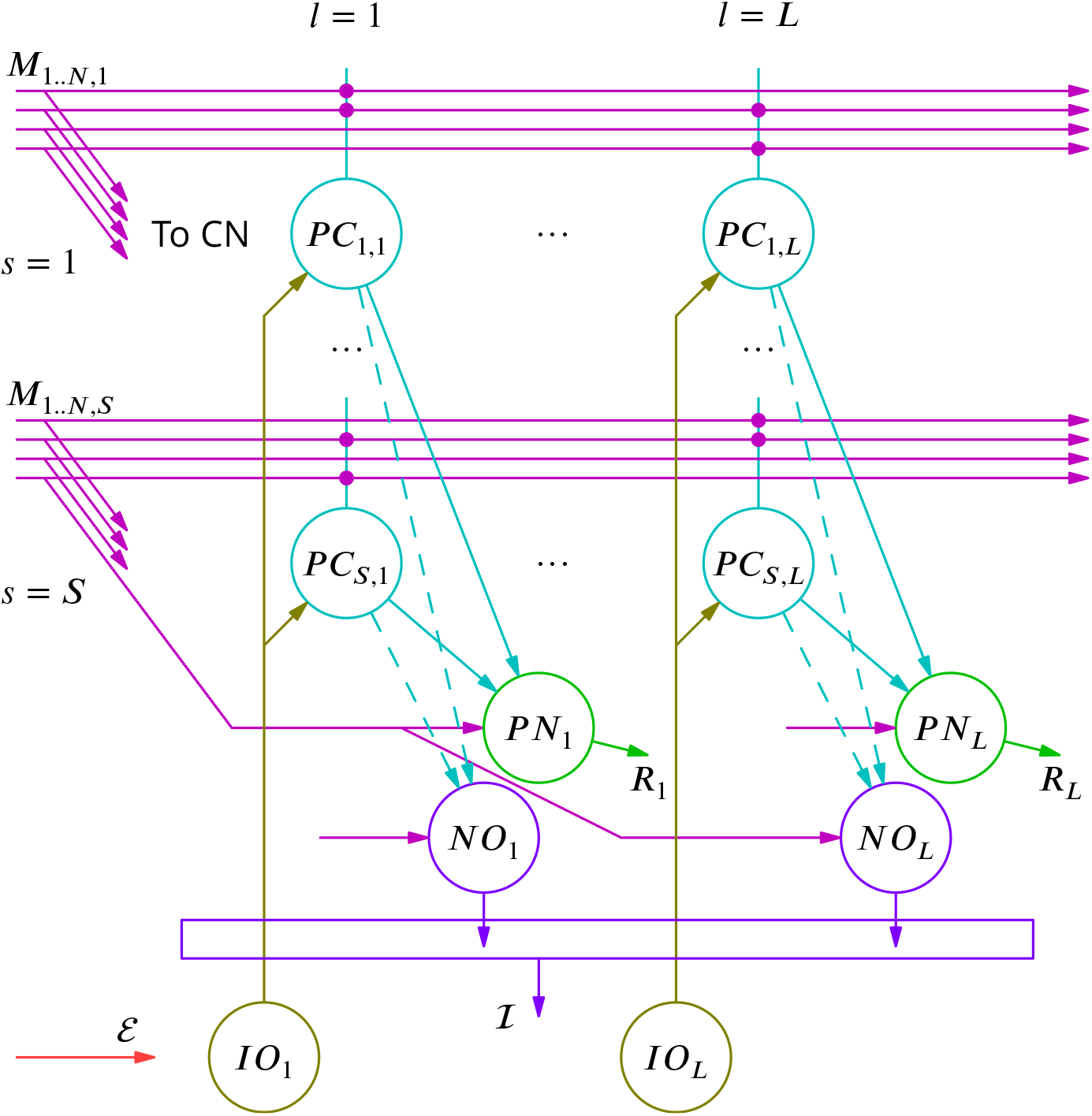
Diagram of the simulated network model. The details are explained in the text.

The connectivity was chosen such that all Purkinje cells in the sagittal ‘column’ *l* projected to the *l*-th nuclear projection neurone with identical inhibitory (negative) weights. Mossy fibres were chosen to project to Purkinje cells with groups of *N* fibres at a given sagittal position *s*, contacting the row of *L* Purkinje cells at the sagittal position with probability 1/2. Purkinje cells were considered not to influence nucleo-olivary neurones except during evaluation of movement errors and the induction of plasticity (described below). The activity of Purkinje and cerebellar nuclear cells in the absence of climbing fibre activity was thus

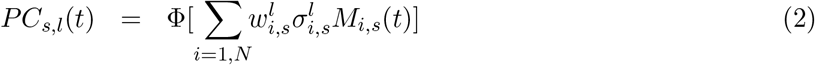

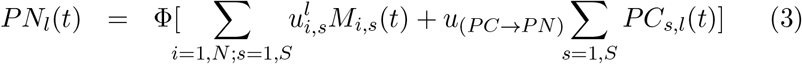

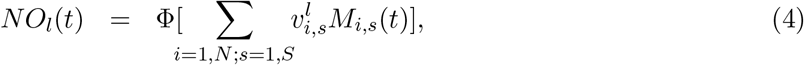

where 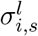 enforces the 1/2 probability of connection between a Purkinje cell and a parallel fibre that traverses its dendritic tree: 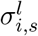 is equal to 1 or 0 with probability 1/2, independently drawn for each triple (*i, s, l*). The *f* − *I* curve Φ was taken to be a saturating threshold linear function, {Φ(*r*) = 0 for *r* < 0, Φ(*I*) = *r*, for 0 < *r* < *r_max_* and Φ(*I*) = *r_max_* for *r* > *r_max_*}. The weights 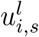 were non-plastic and chosen such that a given mossy fibre (*i, s*) contacted a single projection neurone and a single nucleo-olivary neurone among the *L* possible ones of each type. In other words, for each mossy fibre index (*i, s*), a number *l*_(*i,s*)_ was chosen at random with uniform probability among the *L* numbers {1, ⋯, *L*} and the weights 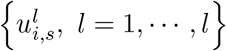 were determined as

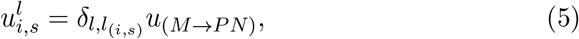

where *u*_(*M*→*PN*)_ was a constant. The weights 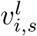 and 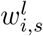 were plastic and followed the learning dynamics described below (see Eq. 11 and 12).

The learning task itself consisted of producing, in response to *μ* = 1, ⋯, *p*, spatiotemporal patterns of mossy fibre inputs 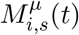, the corresponding output target rates of the projection neurones 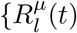. For each pattern *μ*, the inputs were obtained by choosing at random with uniform probability *NS*/2 active fibres. For each active fibre (*i, s*), a time bin 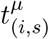 was chosen at random with uniform probability and the activity of the fibre was set to 1 in this time bin, 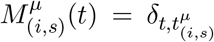. The activity was set to zero in all time bins for the *NS*/2 inactive fibres. The target rates where independently chosen with uniform probability between 0 and 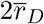 for each projection neurone in each pattern *μ*, where 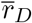 is the desired average firing rate for both projection and nucleo-olivary neurones in the cerebellar nuclei.

The olivary neurones were not explicitly represented. It was assumed that the *L* × *S* Purkinje cells were contacted by *L* climbing fibres with one climbing fibre contacting the *S* Purkinje cells at a given lateral position.

The learning algorithm then proceeded as follows. Patterns 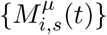 were presented sequentially. After pattern *μ* was chosen, perturbations of Purkinje cell firing by complex spikes were generated as follows. The probability that each climbing fibre emitted a perturbation complex spike was taken to be *ρ* per pattern presentation; when a climbing fibre was active, it was considered to perturb the firing of its Purkinje cells in a single time bin chosen at random. Denoting by *η_l_*(*t*) = 1 that climbing fibre *l* had emitted a spike in time bin *t* (and *η_l_*(*t*) = 0 when there was no spike), the *S* firing rates of the Purkinje cell at position *l* (see Eq. 3) were taken to be

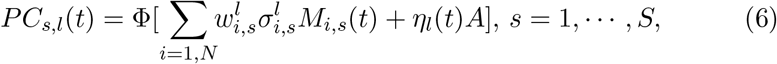

where *A* defines the amplitude of the complex-spike perturbation of Purkinje cell firing.

Given the pattern *μ* and the firing of Purkinje cells (Eq. (6)), the activities of cerebellar nuclear neurones were given by Eqs. (3) and (4). The current ‘error’ for pattern/movement *μ* was quantified by the distance of the projection neurones’ activity from their target rates

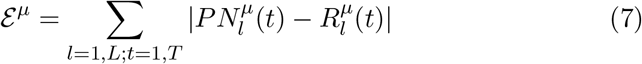

The learning step after the presentation of pattern *μ* was determined by the comparison (not explicitly implemented) in the olivary neurones between the excitation 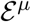 and the inhibition 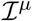 coming from the discharges of nucleo-olivary neurones with

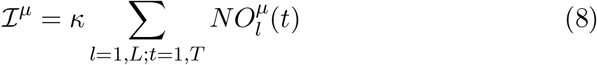

where *κ* ≈ 0.87 was a constant chosen empirically in order to reduce the initial imbalance between error and inhibition in the olive. An error complex spike was propagated to all Purkinje cells after a ‘movement’ when the olivary activity 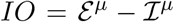 was positive. Accordingly, modifications of the weights of mossy fibre synapses on perturbed Purkinje cells (*w*) and on nucleo-olivary neurones (*v*) were determined after presentation of pattern *μ* by the sign c of *IO, c* = sign(*IO*), as,

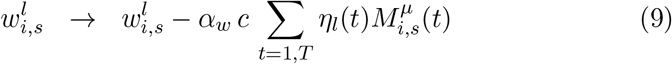

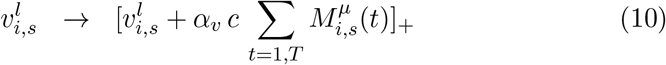

where the brackets served to enforce a positivity constraint on the weights 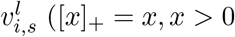 and [*x*]_+_ = 0, *x* < 0).

In the reported simulations, the initial weights of the plastic synapses were drawn from uniform distributions. For *M* → *PC* synapses 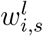 was drawn from 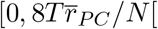 and for *M* → *NO* synapses, weights 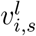 were drawn from 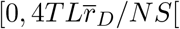. The non-plastic weights in Eq. (3) and (5) were identical and constant for all synapses of a given type

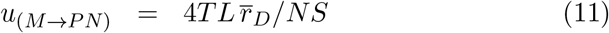

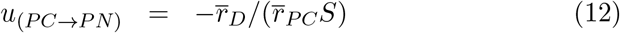

These initial weights ensured initial average firing rates close to 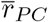 and 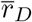 in Purkinje cells and the cerebellar nuclear neurones, respectively.

The parameters used in the reported simulation are provided in Table 1.

**Table 1:**
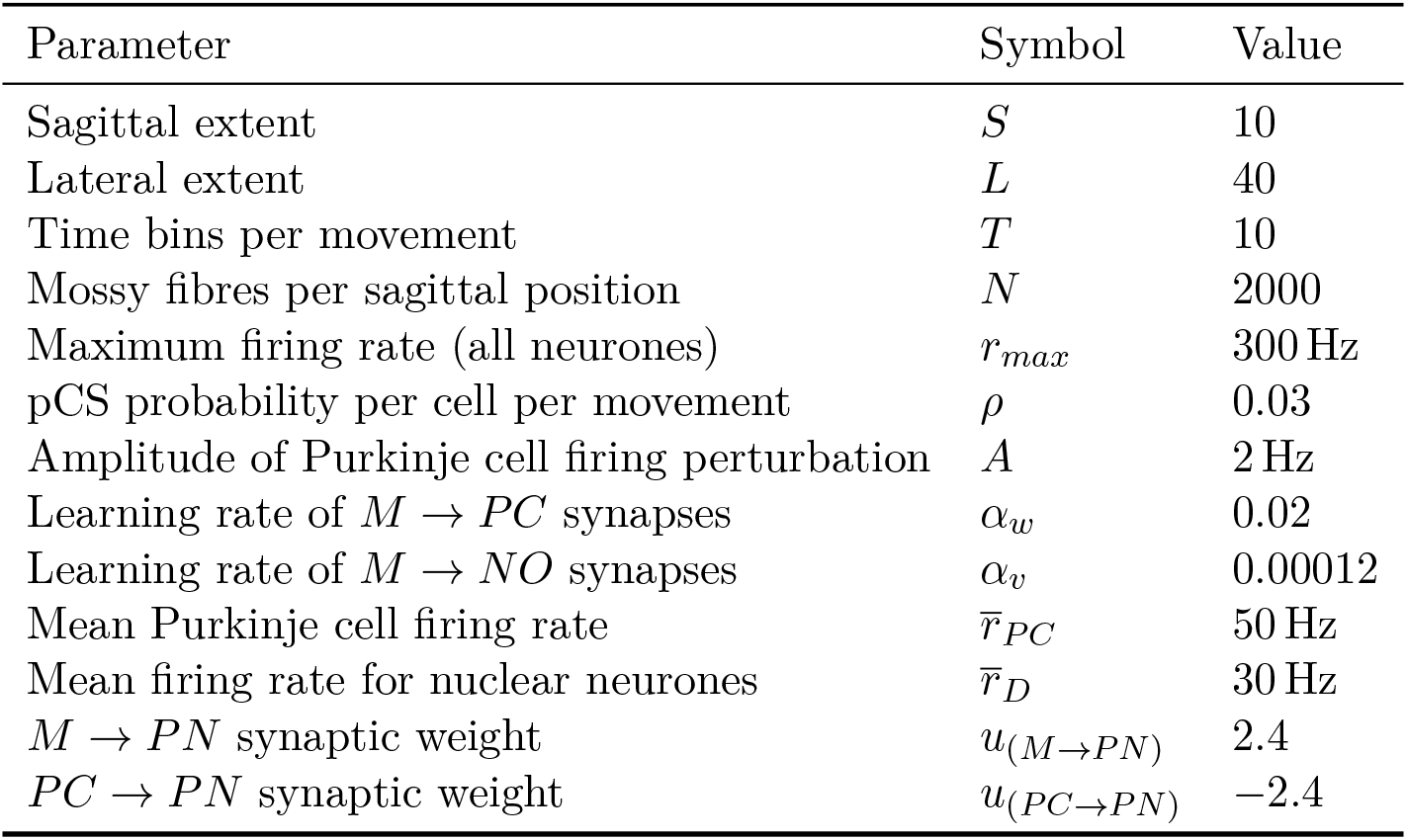
Parameters used in the simulation shown in Fig. 2 and §7

Averaged over two patterns, the initial firing rates were 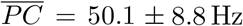 and 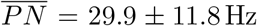 (mean ± sd of *n* = *pLT* = 800 rates for cerebellar nuclear neurones and *n* = *pLTS* = 8000 rates for Purkinje cells).

Analysis in section §8 will show that the four key parameters governing convergence of our learning algorithm are the learning rates of mossy fibre–Purkinje cell synapses *α_w_* and mossy fibre–nucleo-olivary neurone synapses *α_v_*, as well as the probability of a perturbation complex spike occurring in a given movement in a given cell *ρ*, and the resulting amplitude of the perturbation of Purkinje cell firing *A*. Increasing each of these 10 % individually increased the final error (averaged over trials 70000–75000) by at most 11.6%, indicating that this final output was not ill-conditioned or finely tuned with respect to these parameters.

The simulation was coded in C++.

## 3 Requirements for a general cerebellar learning algorithm

We begin by analysing the current consensus theory of cerebellar learning and highlight some of its potential deficiencies. The learning framework is the following. Execution of a movement involves Purkinje cells producing trains of action potentials of varying frequencies at given times (Thach, 1968). We consider only the rate *r*(*t*), which is a function of time in the movement. Purkinje cell output is combined with direct mossy fibre input in the cerebellar nuclei, whose neurones must also produce varying spike trains. Purkinje cells must learn to adapt the output of their cerebellar nuclear neurone targets so as to minimise the movement error. Mossy fibre firing varies during a movement, but we assume for simplicity that it does not change during learning. The minimisation of error is to be achieved through plasticity of parallel fibre synapses, which drive Purkinje cell activity (possibly with complementary modifications involving molecular layer interneurones; Jörntell and Ekerot, 2002, 2003; Mittmann and Häusser, 2007). We shall assume that LTP of active parallel fibre synapses increases the rate r and LTD decreases it (Lev-Ram et al., 2003).

We plot possible cell- and movement-specific error signals carried by a climbing fibre for two (sub)movements against the firing rate of a Purkinje cell in Fig. 3A. The frequency of complex spikes can be increased or decreased below their baseline rate of about 1 Hz (e.g. Ke et al., 2009). The movements have different optimal frequencies and opposite slopes. We first consider movement *m*_1_, which is associated with a complex spike rate that is positively correlated with *r*. As complex spikes trigger LTD and parallel fibres active in their absence are potentiated, we expect a stable or set point for the rate, which is driven by the plastic synapses. The set point would correspond to zero error and the non-zero baseline rate of complex spikes. Thus, if the Purkinje cell rate is above optimum during *m*_1_, the synapses driving it will be depressed, returning its activity towards the optimum; a converse argument applies if the rate falls below the optimum.

**Figure 3:**
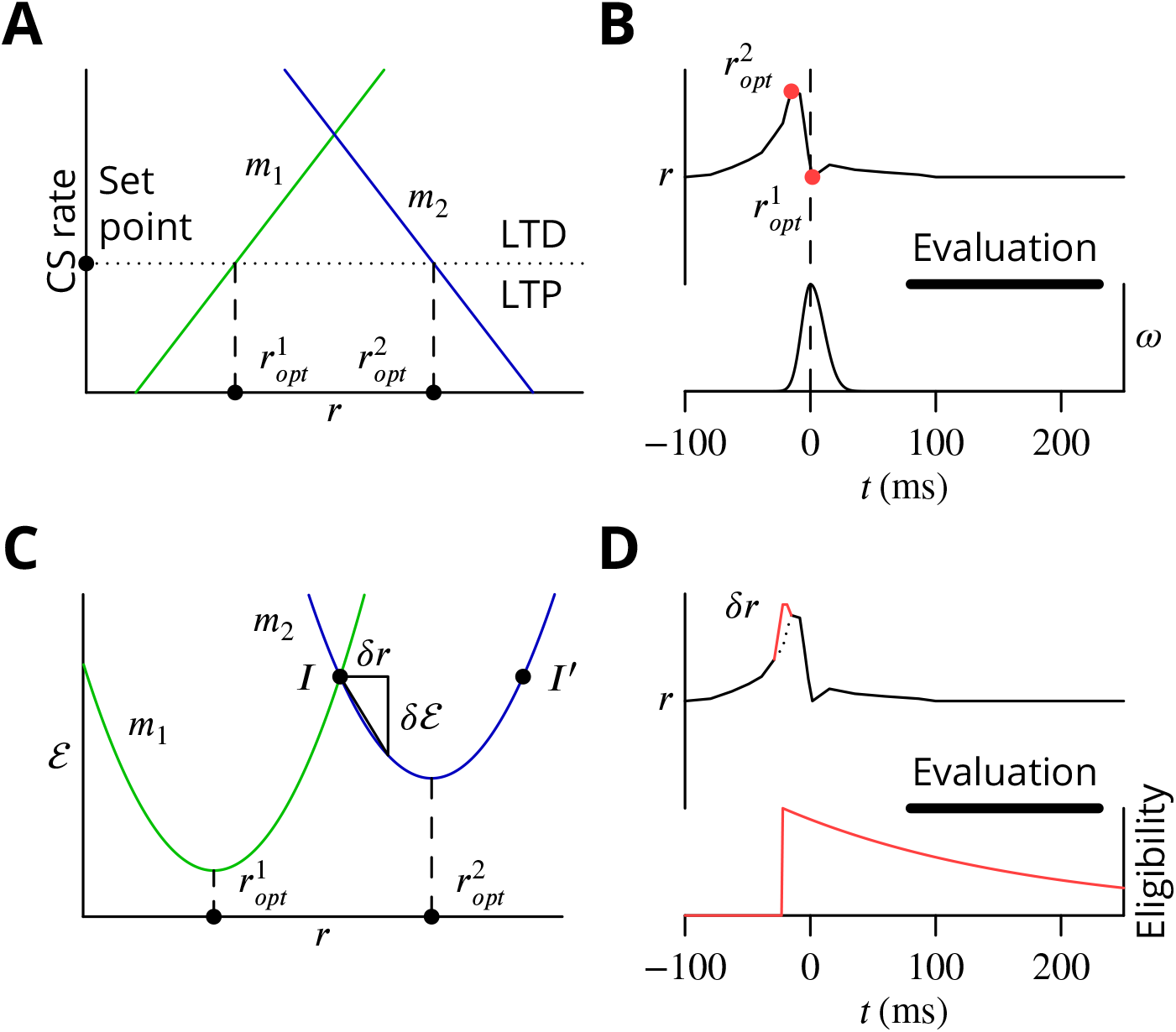
Analysis of requirements for cerebellar learning. **A**. Possible cell- and movement-specific error signals represented by complex spike rate (CS rate) for (sub)movements *m*_1_ and *m*_2_ as functions of Purkinje cell firing rate (*r*, optima *r_opt_*) within the consensus learning theory. The CS rate can instruct the direction of plasticity. **B**. Illustration of firing and movement time courses during cerebellar control of saccades. Eye movement (*ω*, angular velocity, bottom-right axis) is preceded by a short burst of increased activity (*r*, top and left axis) in Purkinje cells. Different time points have different optima (*r_opt_*, red dots). The accuracy of the saccade can only be evaluated once it is complete and visual information has been processed (Evaluation, thick black line). **C**. Possible global error function 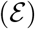 whose value cannot directly instruct optimising plasticity. Thus, points *I* and *I*′ have the same error, but require plasticity of different signs to approach 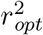. Stochastic gradient descent involves perturbing the rate (*δr*) and determining the resulting error change 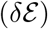 to obtain the gradient. **D**. Stochastic gradient descent illustrated in the context of a saccade. A perturbation of the firing rate (*δr*, red vs. dotted) tags simultaneously active synapses in that cell (eligibility trace at bottom, red), allowing synapse-specific plasticity according to the movement evaluation.

This leads us to a first constraint on climbing fibre activity in the current theory. The optimum rate of the Purkinje cell is defined by the set point of climbing fibre activity. Because coordinated movements require parallel output from many parts of the cerebellum, it is unavoidable that individual Purkinje cells contribute to the control of many different movements with different optima. The error processing that generates climbing fibre activity would therefore need to be endowed with sufficient information to define this set point correctly for all cell and movement pairs.

Movement *m*_2_ has a negative slope. We see that applying the same plasticity rule will result in unstable learning, causing the firing rate to diverge from the optimum. Thus, if the rate is too low, the increased error will cause a high rate of complex spikes, leading to depression of the synapses driving the Purkinje cell, which will then fire at an even lower frequency, and so on. We therefore identify a second constraint: increases of Purkinje cell firing must always affect movements so as to increase complex spike frequency 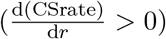. The same must be true within the circuitry of the cerebellum. The synaptic signs of the Purkinje cell–nucleo-olivary neurone–inferior olive loop do appear to satisfy this condition (Chaumont et al., 2013).

Although these requirements on the relations between error, complex spike and Purkinje cell firing appear near-impossible to satisfy in general for arbitrary movements, it may be possible to do so partially in the context of some constrained behaviours, notably eye movements and simple reflexes. These provide some of the best studied models of cerebellar learning: the vestibulo-ocular reflex (Robinson, 1976; Ito et al., 1974; Blazquez et al., 2004), nictitating membrane response/eyeblink conditioning (McCormick et al., 1982; Yeo et al., 1984; Yeo and Hesslow, 1998), saccade adaptation (Optican and Robinson, 1980; Dash and Thier, 2014; Soetedjo et al., 2008) and regulation of limb movements by withdrawal reflexes (Ekerot et al., 1995; Garwicz et al., 2002). All of these motor behaviours have in common that there could conceivably be a fixed mapping between an error and a suitable corrective action. Thus, the compensatory eye movement is exactly determined by the retinal slip, while a puff of air directed at the cornea should always trigger a blink protecting the eye. Such fixed error-correction relations may have been exploited during evolution to create optimised correction circuitry.

The temporal characteristics of cerebellar motor control tasks nevertheless create additional obstacles for the traditional learning algorithm, because a single evaluation of the result of a movement may have to specify complex temporal activity sequences. This can be seen in the context of saccade adaptation. The Purkinje cells required for saccade adaptation display a characteristic population activity burst during a saccade (Thier et al., 2000); this is diagrammed in Fig. 3B (adapted from Herzfeld et al., 2015). This burst is quite brief, of the order of 50 ms. During the burst, Purkinje cells display abrupt changes of firing frequency, with changes of 100 Hz in 20 ms often being observed. Each time point of the population burst can be considered as a distinct optimal frequency that must be learnt. The rapid modulation of Purkinje cell firing therefore implies that very different plasticity occurs over intervals of no more than 20 ms.

But the accuracy of a saccade can only be evaluated once it is complete. Vision is moreover a particularly slow sense. This implies that specific and different plasticity outcomes at time points 20 ms apart during the Purkinje cell population burst must all be determined by a unique evaluation occurring some time later. The timing of that evaluation and the complex spikes it generates may be subject to significant jitter. The window for evaluation (based upon the post-movement period when excess complex spikes are observed; Soetedjo et al., 2008) shown in Fig. 3B,D illustrates the problem—it is broader than the saccade-related activity and movement. Even more challenging, the burst of simple spike activity shown would presumably require LTP to become established, implying an absence of complex spikes, as can be observed throughout the evaluation window under the appropriate error conditions (Soetedjo et al., 2008). In other words, the absence of complex spikes within a window of 150 ms would need to specify strong LTP at a time point 100 ms before the beginning of the evaluation window, but no plasticity at 80 ms before the window^2^.

Although it may not be impossible for the traditional learning algorithm to overcome these obstacles in the case of constrained movements like those of the eye, it seems extremely challenging. And implementing such an algorithm capable of solving more complex, arbitrary and high-dimensional motor control tasks would pose even greater difficulties.

We next analyse the requirements for a more general cerebellar learning algorithm. More realistic, less-informative global error functions are diagrammed in Fig. 3C. The global error 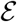 for two movements (or submovements) is shown as a function of the firing rate *r* of the Purkinje cell, assuming nothing else changes in the network (including the mossy fibre input). Although the parabolic forms are of course arbitrary, learning should seek whatever neighbouring minima exist in the error functions, such as 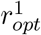 and 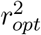. Contrary to the functions in Fig. 3A, those of Fig. 3C are not instructive of the sign of plasticity required to approach the optimum. Thus, neither the existence nor the absolute value of the error signal is sufficient to indicate the required sign of plasticity. This can be seen by considering points *I* and *I*′ on the error function of *m*_2_. They have identical error intensities, but from *I* an increase in r and therefore LTP is required, while from *I*′ a decrease and therefore LTD is needed.

A key constraint on the learning algorithm is that the coordinates (*r* and 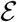) of the error function minima can vary between movements and even with time within movements. That the *r_opt_* can vary within and between movements is uncontroversial. If we imagine for the sake of argument that the firing of a Purkinje cell controls the force of a muscle^3^, different motor states may require different degrees of contraction, implying different Purkinje cell firing rates. In contrast, allowing for variation of the minimum error between movements may initially appear unnecessary. However, the error in a movement depends upon the degree of optimisation throughout the whole network, which may be substantially different for the two movements, independently of the behaviour of the cell depicted. We recall that the curves of Fig. 3C show the error as the rate of that individual cell is altered.

The above analysis strongly suggests that it is impractical for the cerebellar circuitry to deduce optimal network changes from the movement error alone. At the very least, the sign of the gradient of the error function must be determined if plasticity is to reduce the error reliably and ultimately minimise it.

The decorrelation theory (Dean et al., 2002; Dean and Porrill, 2014) offers an elegant implicit gradient descent algorithm, but we believe that the detailed implementation suggested (Menzies et al., 2010) is unable to solve the temporal credit assignment problem arising in movements that can only be evaluated upon completion. We shall also see below that the present implementation employs different plasticity rules to those of the theory we shall develop.

Stochastic gradient descent (Minsky, 1961) provides a plausible algorithm for solving the cerebellar learning problem via gradient descent. This algorithm involves a sequence of small trial network modifications, with consolidation of those that improve the behaviour. Applied here (Fig. 3C), a perturbation *δr* is effected and the resulting change of error 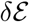 determined. Knowledge of the signs of both *δr* and 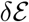 is necessary and sufficient information to minimise the error. Assuming the perturbation is sparse within the network, it can be used to tag the perturbed circuit elements, creating an eligibility trace that can be exploited to guide plasticity specific to those elements (Fig. 3D).

## 4 The complex spike as trial and error

Our attempt to design a cerebellar implementation of stochastic gradient descent began with a search for a source of perturbation *δr*. The fact that Purkinje cells can contribute to different movements with arbitrary and unknown sequencing imposes an implementation constraint preventing simple-minded approaches like comparing movements performed twice in succession. We recall that we assume that no explicit information categorising or identifying movements is available to the Purkinje cell. It is therefore necessary that knowledge of both the presence and sign of *δr* be available within the context of a single movement execution.

In practice, a number of different perturbation mechanisms can still satisfy these requirements. For instance, any binary signal would be suitable, since the sign of the perturbation with respect to its mean would be determined by the simple presence or absence of the signal. Several plausible mechanisms along these lines have been proposed, including external modulatory inputs (Doya and Sejnowski, 1988; Fiete et al., 2007), failures and successes of synaptic transmission (Seung, 2003) or the absence and presence of action potentials (Xie and Seung, 2004). However, none of these mechanisms has yet attracted experimental support at the cellular level.

In the cerebellar context, parallel fibre synaptic inputs are so numerous that the correlation between individual input variations and motor errors is likely to be extremely weak, whereas we seek a perturbation that is sufficiently salient to influence ongoing movement. Purkinje cell action potentials are also a poor candidate, because they are not back-propagated to parallel fibre synapses (Stuart and Häusser, 1994) and therefore probably cannot guide their plasticity, but the ability to establish a synaptic eligibility trace is required. We considered bistable firing behaviour of Purkinje cells (Loewenstein et al., 2005; Yartsev et al., 2009), with the down-state (or long pauses) representing a clear perturbation towards lower (zero) firing rates. However, exploratory plasticity experiments did not support this hypothesis and the existence of bistability in vivo is disputed (Schonewille et al., 2006a).

We then considered another possible perturbation of Purkinje cell firing: the complex spike triggered by the climbing fibre. We note that there are probably two types of inferior olivary activity. Olivary neurones mediate classical error signalling triggered by external synaptic input, but they also exhibit continuous and irregular spontaneous activity in the absence of overt errors. We suggest the spontaneous climbing fibre activations cause synchronised *perturbation complex spikes* (pCSs) in small groups of Purkinje cells via the ~ 1:10 inferior olivary–Purkinje cell divergence (Schild, 1970; Mlonyeni, 1973; Caddy and Biscoe, 1976) and dynamic synchronisation of olivary neurones (through electrical coupling Llinás and Yarom, 1986; Bazzigaluppi et al., 2012a, or common drive). The excitatory perturbation—a brief increase of firing rate (Ito and Simpson, 1971; Campbell and Hesslow, 1986; Khaliq and Raman, 2005; Monsivais et al., 2005)—feeds through the cerebellar nuclei (changing sign; Bengtsson et al., 2011) to the ongoing motor command and causes a perturbation of the movement, which in turn may modify the error of the movement. Harris (1998) has previously proposed a cerebellar learning algorithm that involves a similar perturbation role for complex spikes, although the rest of our implementation differs from his.

The perturbations could guide learning in the following manner. If a perturbation complex spike results in an increase of the error, the raised activity of the perturbed Purkinje cells was a mistake and reduced activity would be preferable; parallel fibre synapses active at the time of the perturbing complex spikes should therefore be depressed. Conversely, if the perturbation leads to a reduction of error (or does not increase it), the increased firing rate should be consolidated by potentiation of the simultaneously active parallel fibres.

How could an increase of the error following a perturbation be signalled to the Purkinje cell? We suggest that the climbing fibre also performs this function, although we postpone the description of a mechanism for achieving it until later. Specifically, if the perturbation complex spike increases the movement error, a secondary *error complex spike* (eCS) is emitted shortly afterwards, on a time scale of the order of 100 ms. This time scale is assumed because it corresponds to the classical error signalling function of the climbing fibre, because it allows sufficient time for feedback via the error modalities known to elicit complex spikes (touch, pain, balance, vision) and because such intervals are known to be effective in plasticity protocols (Wang et al., 2000; Sarkisov and Wang, 2008; Safo and Regehr, 2008). The interval could also be influenced by the oscillatory properties of olivary neurones (Llinás and Yarom, 1986; Bazzigaluppi et al., 2012b).

The predicted plasticity rule is therefore as diagrammed in Fig. 4. Only granule cell synapses active simultaneously with the perturbation complex spike undergo plasticity (Fig. 4A,B), with the sign of the plasticity being determined by the presence or absence of a subsequent error complex spike. Granule cell synapses active in the absence of a synchronous perturbation complex spike should not undergo plasticity, even if succeeded by an error complex spike (Fig. 4C,D). We refer to these different protocols with the abbreviations (and give our predicted outcome in parenthesis): G_ _ (no change), GP_ (LTP), G_E (no change), GPE (LTD), where G indicates granule cell activity, P the presence of a perturbation complex spike and E the presence of an error complex spike. Note that both granule cells and climbing fibres are likely to be active in high-frequency bursts rather than the single activations idealised in Fig. 4.

**Figure 4:**
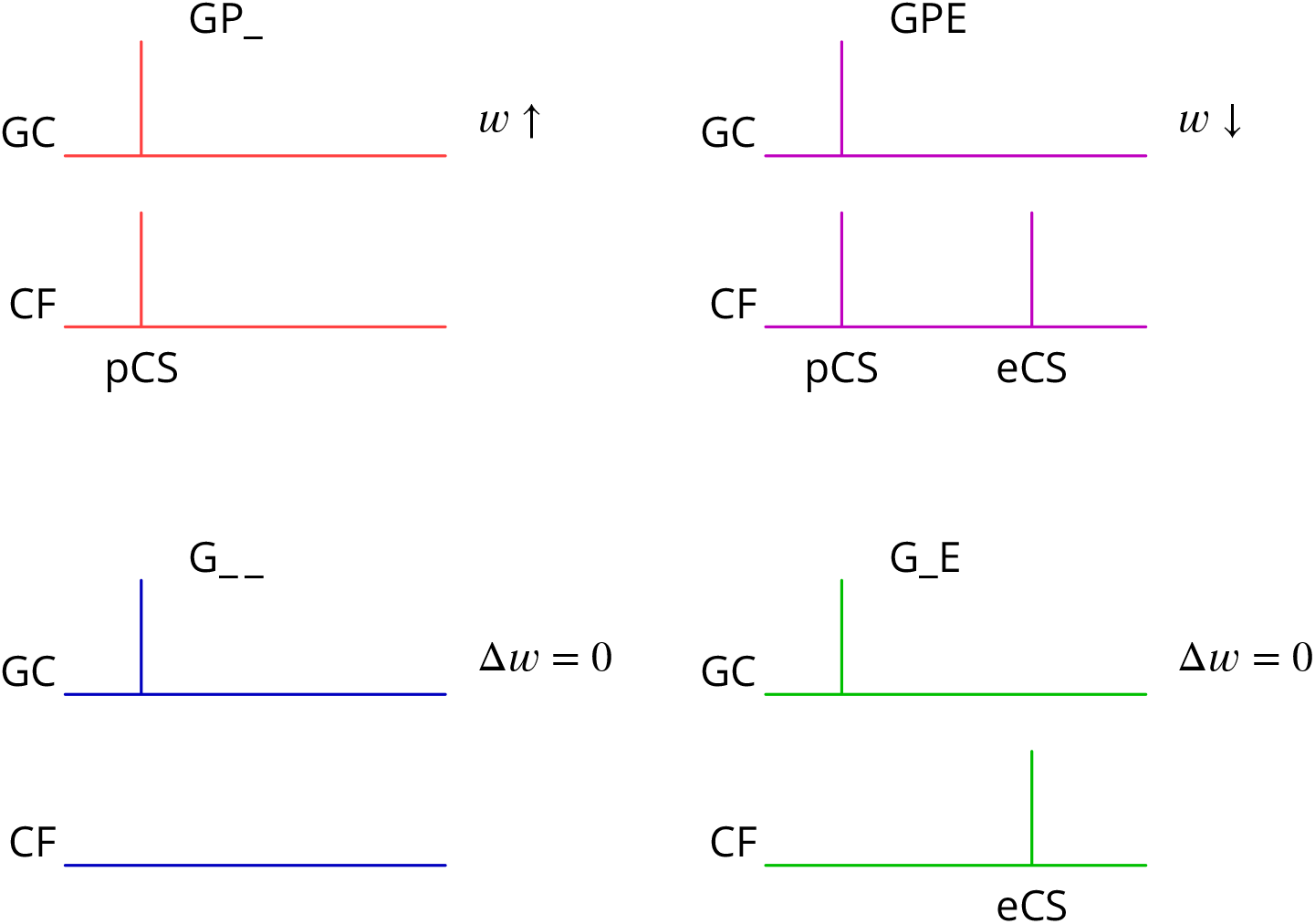
Predicted plasticity rules. Synchronous activation of granule cell synapses and a perturbation complex spike (pCS) leads to LTP (GP_, increased synaptic weight *w*; top left, red), while the addition of a succeeding error complex spike (eCS) leads to LTD (GPE, top right, magenta). The bottom row illustrates the corresponding ‘control’ cases from which the perturbation complex spike is absent; no plasticity should result (G_ _ blue and G_E green).

Several of the predictions of this rule appear to be incompatible with the current consensus. Thus, parallel fibre synapses whose activity is simultaneous with (GP_) or followed by a complex spike (G_E) have been reported to be depressed (Sakurai, 1987; Crepel and Jaillard, 1991; Lev-Ram et al., 1995; Coesmans et al., 2004; Safo and Regehr, 2008; Gutierrez-Castellanos et al., 2017), while we predict potentiation and no change, respectively. Furthermore, parallel fibre activity alone (G_ _) leads to potentiation (Lev-Ram et al., 2002; Jörntell and Ekerot, 2002; Lev-Ram et al., 2003; Coesmans et al., 2004; Gutierrez-Castellanos et al., 2017), while we predict no change.

## 5 Synaptic plasticity under physiological conditions

As described above, the plasticity rules we predict for parallel fibre–Purkinje cell synapses are, superficially at least, close to the opposite of the consensus in the literature, to which we have contributed (Bidoret et al., 2009; Ly et al., 2013; Bouvier et al., 2016). Current understanding of the conditions for inducing plasticity gives a key role to the intracellular calcium concentration (combined with nitric oxide signalling; Coesmans et al., 2004; Bouvier et al., 2016), whereby high intracellular calcium concentrations are required for LTD and moderate concentrations lead to LTP. Standard experimental conditions for studying plasticity in vitro, notably the extracellular concentration of calcium, are likely to result in more elevated intracellular calcium concentrations during induction than pertain physiologically. Recognising that this could alter plasticity outcomes, we set out to test whether our predicted plasticity rules might be verified under more physiological conditions.

We made several changes to standard protocols (see Methods); one of the changes was cerebellum-specific, but the others also apply to in vitro plasticity studies in other brain regions. We did not block GABAergic inhibition. We lowered the extracellular calcium concentration from the standard 2 mM (or higher) used in slice work to 1.5 mM, which is near the maximum values measured in vivo in rodents (Jones and Keep, 1988; Silver and Erecińska, 1990). We only used weak granule cell layer stimuli, which results in sparse and spatially dispersed parallel fibre activity, avoiding the compact bundles of parallel fibres almost universally employed. Interestingly, it has been reported that standard protocols using granule cell stimulation are unable to induce LTD (Marcaggi and Attwell, 2007). We used a pipette solution designed to prolong energy supply in extended cells like the Purkinje cell (see Methods). Experiments were carried out in adult mouse sagittal cerebellar slices using otherwise standard patch-clamp techniques. The use of this combination of physiological conditions for in vitro plasticity studies has not been previously reported for the cerebellum, indeed for any brain region, with the notable exception of work from the group of Indira Raman (e.g. Pugh and Raman, 2008).

During induction, performed in current clamp without any injected current, the granule cell input consisted of a burst of five stimuli at 200 Hz, reproducing the propensity of granule cells to burst at high frequencies (Chadderton et al., 2004; Jörntell and Ekerot, 2006). The climbing fibre input reflected the fact that these can occur in very high-frequency bursts (Eccles et al., 1966; Maruta et al., 2007). We used two stimuli at 400 Hz to represent the perturbation complex spike and four for the subsequent error complex spike if it was included in the protocol. In a fraction of cells, the climbing fibre stimuli after the first were not reliable; our grounds for including these cells are detailed in the Methods. The interval between the two bursts of climbing fibre stimuli when the error complex spike was present was about 100 ms. We increased the interval between induction episodes from the standard one second to two, to reduce any accumulating signal during induction. 300 such induction sequences were applied (Lev-Ram et al., 2002).

Pairs of granule cell test stimuli with an interval of 50 ms were applied at 0.1 Hz before and after induction; EPSCs were recorded in voltage clamp at −70 mV. Pairs of climbing fibre stimuli with a 2.5 ms interval were applied at 0.5 Hz throughout the test and induction periods; the interleaved test granule cell stimulations were sequenced 0.5 s before the climbing fibre stimulations. During induction, granule cell stimuli were applied in phase or in anti-phase with these perturbation signals as required for the different protocols described below. The analysis inclusion criteria and amplitude measurement for the EPSCs are detailed in the Methods. The average amplitude of the granule cell EPSCs retained for analysis was −62 ± 46 pA (mean ± s.d., *n* = 58). The rise and decay time constants (of the global averages) were 0.74 ± 0.36ms and 7.2 ± 2.7ms (mean ± sd), respectively.

We first show the protocols relating to LTP (Fig. 5). A granule cell burst was followed by a distant perturbation climbing fibre stimulus or the two inputs were activated simultaneously. In the examples shown, the protocol with simultaneous activation (GP_, Fig. 5C,D) caused a potentiation of about 40 %, while the temporally separate inputs caused a smaller change of 15 % in the opposite direction (G_ _, Fig. 5A,B). We note that individual outcomes were quite variable; group data and statistics will be shown below. The mean paired-pulse ratio in our recordings was A2/A1 = 1.75 ± 0.32 (mean ± sd, *n* = 58). As here, no significant differences of paired-pulse ratio were observed with any of the plasticity protocols: plasticity – baseline difference for GP_, mean −0.08, 95 % confidence interval (−0.34, 0.20), *n* = 15; GPE, mean 0.12, 95% c.i. (−0.07, 0.33), *n* = 10; G_ _, mean −0.01, 95% c.i. (−0.24, 0.24), *n* = 18; G_E, mean −0.09, 95% c.i. (−0.23, 0.29), *n* = 15.

**Figure 5:**
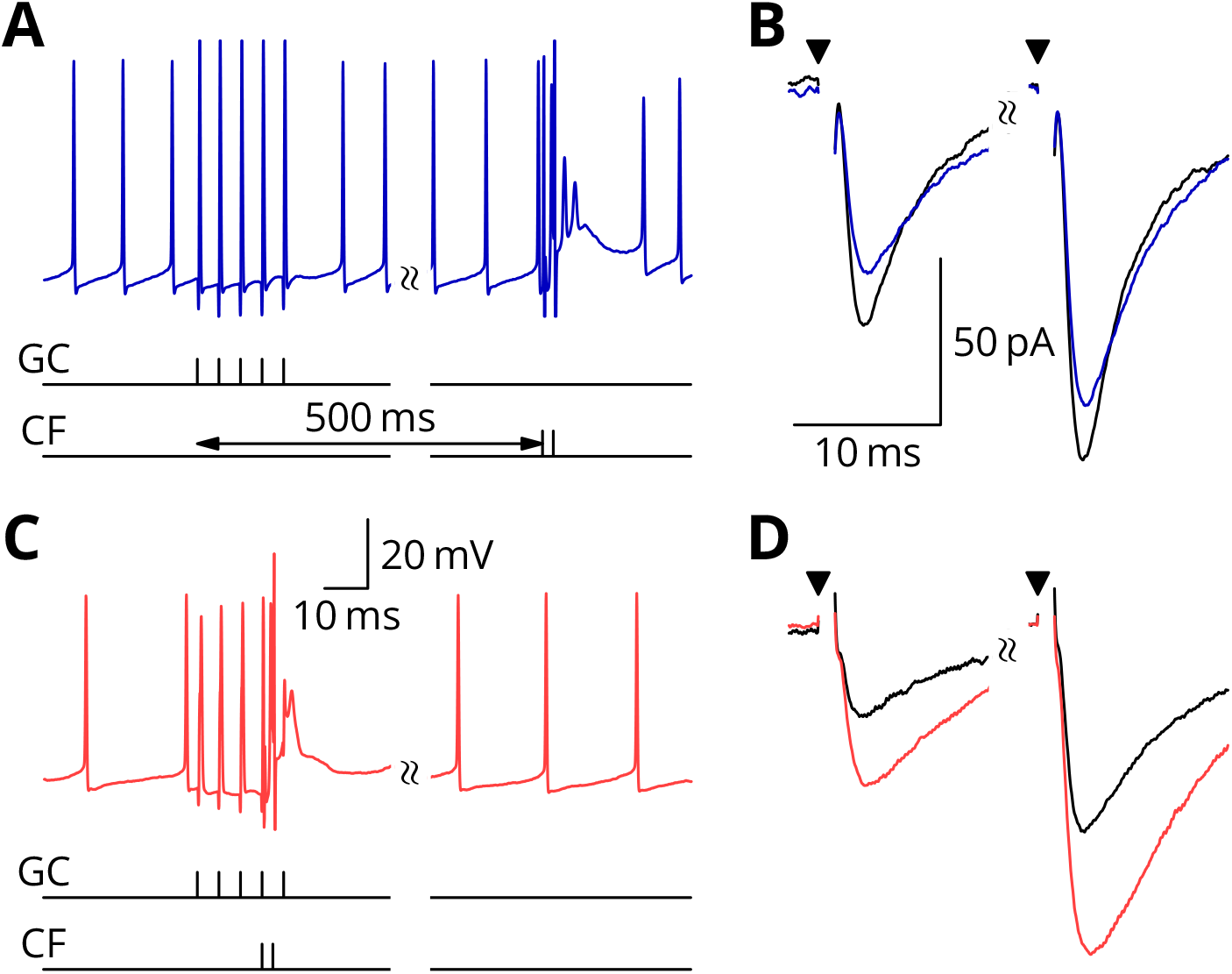
Simultaneous granule cell and climbing fibre activity induces LTP. **A**. Membrane potential (blue) of a Purkinje cell during an induction protocol (G_ _) where a burst of 5 granule cell stimuli at 200 Hz was followed after 0.5 s by a pair of climbing fibre stimuli at 400 Hz. **B**. Average EPSCs recorded up to 10 minutes before (black) and 20–30 minutes after the end of the protocol of A (blue). Paired test stimuli (triangles) were separated by 50 ms and revealed the facilitation typical of the granule cell input to Purkinje cells. In this case, the induction protocol resulted in a small reduction (blue vs. black) of the amplitude of responses to both pulses. **C**. Purkinje cell membrane potential (red) during a protocol (GP_) where the granule cells and climbing fibres were activated simultaneously, with timing otherwise identical to A. **D**. EPSCs recorded before (black) and after (red) the protocol in C. A clear potentiation was observed in both of the paired-pulse responses.

During induction, cells would generally begin in a tonic firing mode, but nearly all ceased firing by the end of the protocol. Specimen sweeps are shown in Fig. 6 from later during induction protocols, when spiking had ceased. The protocol was predicted to produce LTD. As before, a granule cell burst was paired with the perturbation climbing fibre, but now a longer burst of climbing fibre stimuli was appended 100 ms later, representing an error complex spike (GPE, Fig. 6C,D). A clear LTD of about 40% developed following the induction. In contrast, if the perturbation complex spike was omitted, leaving the error complex spike (G_E, Fig. 6A,B), no clear change of synaptic weight occurred (an increase of about 10%).

**Figure 6:**
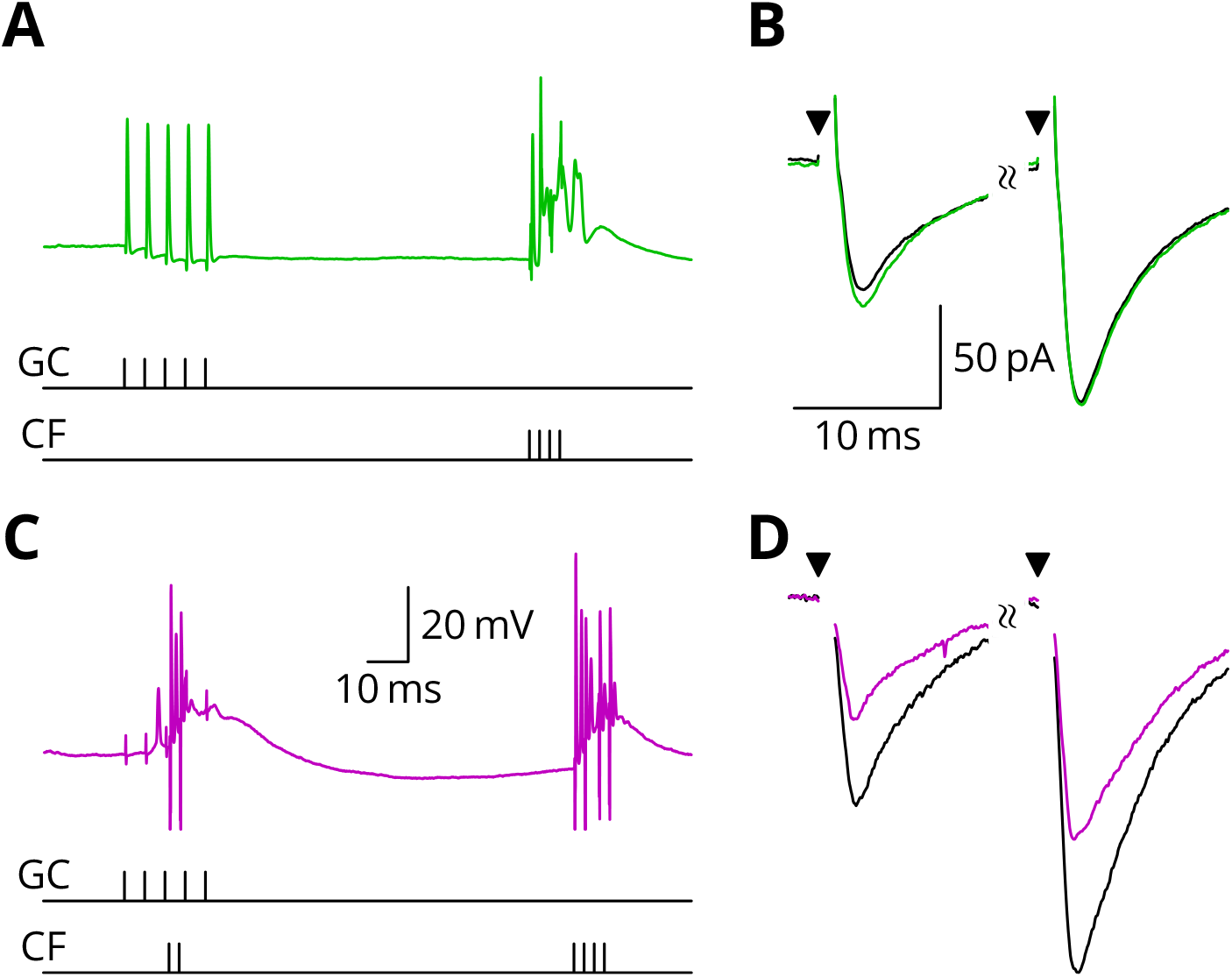
LTD requires simultaneous granule cell and climbing fibre activity closely followed by an additional complex spike. **A**. Membrane potential of a Purkinje cell (green) during a protocol where a burst of 5 granule cell stimuli at 200 Hz was followed after 100 ms by 4 climbing fibre stimuli at 400 Hz (G_E). **B**. Average EPSCs recorded up to 10 minutes before (black) and 20–30 minutes after the end of the protocol of A (green). The interval between the paired test stimuli (triangles) was 50 ms. The induction protocol resulted in little change (green vs. black) of the amplitude of either pulse. **C**. Purkinje cell membrane potential (magenta) during the same protocol as in A with the addition of a pair of climbing fibre stimuli simultaneous with the granule cell stimuli (GPE). **D**. EPSCs recorded before (black) and after (magenta) the protocol in C. A clear depression was observed.

The time course of the changes of EPSC amplitude are shown in normalised form in Fig. 7 (see Methods for selection and normalisation procedures). The individual data points of the relative EPSC amplitudes for the different protocols are also shown; they reveal that the data are quite noisy. A numerical summary of the group data and statistical comparisons is given in Table 2.

**Figure 7:**
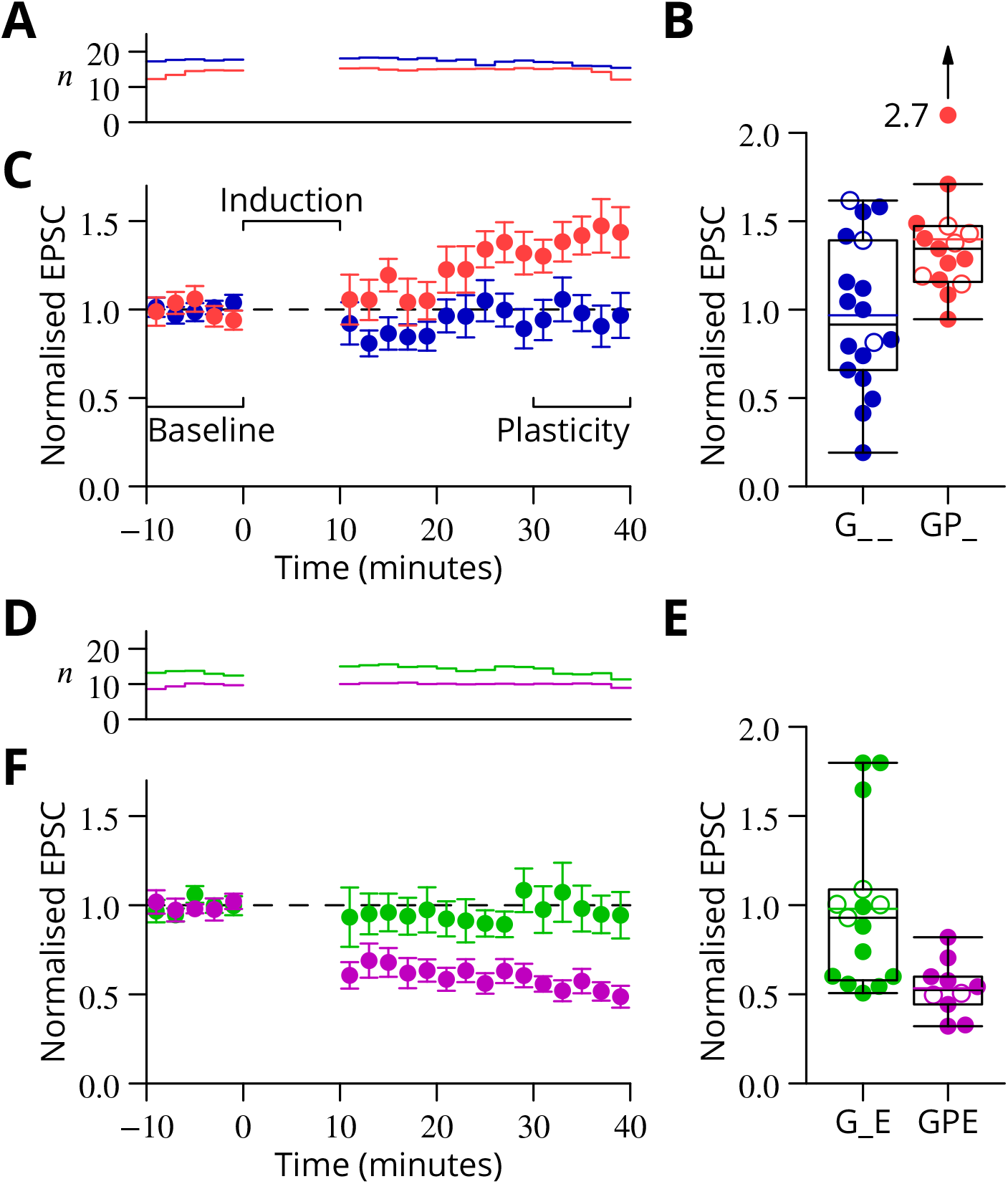
Time course and amplitude of plasticity. **A** number, **B** box-and-whisker plots of individual plasticity ratios (coloured lines represent the means, open symbols represent cells with failures of climbing fibre stimulation; see Methods) and **C** time course of the mean EPSC amplitude for GP_ (red) and G_ _ (blue) protocols of Fig. 5, normalised to the pre-induction amplitude. Averages every 2 minutes, mean ± sem. Non-integer *n* arise because the numbers of responses averaged were normalised by those expected in two minutes, but some responses were excluded (see Methods) and some recordings did not extend to the extremities of the bins. Induction lasted for 10 minutes starting at time 0. **D, E** and **F** Similar plots for the GPE (magenta) and G_E (green) protocols of Fig. 6.

**Table 2:** Group data and statistical tests for plasticity outcomes. In the upper half of the table, the ratios of EPSC amplitudes after/before induction are described and compared with a null hypothesis of no change (ratio = 1). The GP_ and GPE protocols both induced robust changes, while the control protocols (G_ _, G_E) did not. The bottom half of the table analyses differences of those ratios between protocols. The 95 % confidence intervals (c.i.) were calculated using boostrap methods, while the *p*-values were calculated using a two-tailed Wilcoxon rank sum test (see Methods).

| Comparison | Mean | 95 % c.i. | | $p$ | $n$ |
| --- | --- | --- | --- | --- | --- |
| GP_ | 1.40 | 1.27, | 1.71 | 0.0001 | 15 |
| GPE | 0.53 | 0.45, | 0.63 | 0.001 | 10 |
| G__ | 0.97 | 0.78, | 1.16 | 0.77 | 18 |
| G_E | 0.98 | 0.80, | 1.24 | 0.60 | 15 |
| GP_ vs G__ | 0.43 | 0.19, | 0.74 | 0.01 |  |
| GPE vs G_E | -0.44 | -0.71, | -0.24 | 0.0009 |  |
| GP_ vs G_E | 0.42 | 0.14, | 0.73 | 0.005 |  |
| GPE vs G__ | -0.43 | -0.65, | -0.22 | 0.004 |  |
| G__ vs G_E | 0.01 | -0.26, | 0.32 | 0.93 |  |

These results therefore provide experimental support for all four plasticity rules predicted by our proposed mechanism of stochastic gradient descent. We argue in the Discussion that the apparent contradiction of these results with the literature is not unexpected if the likely effects of our altered conditions are considered in the light of known mechanisms of potentiation and depression.

## 6 Extraction of the change of error

Above we have provided experimental evidence in support of the counterintuitive synaptic plasticity rules predicted by our proposed learning mechanism. In that mechanism, following a perturbation complex spike, the sign of plasticity is determined by the absence or presence of a follow-up error complex spike that signals whether the movement error increased (spike present) or decreased (spike absent). We now return to the outstanding problem of finding a mechanism able to extract this *change* of error, 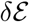.

Several roughly equivalent schemes have been proposed, including subtraction of the average error (Barto et al., 1983) and decorrelation (Dean and Porrill, 2014), a specialisation of the covariance rule (Sejnowski, 1977). Perhaps the most popular method is an elegant implicit extraction of 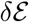 due to Williams (1992). However, in general, these suggestions have not extended to detailed cellular implementations and we believe that designing biologically plausible implementations is not necessarily trivial. For instance, the method of Williams makes use of a small difference between two products, which would require accurate multiplication, an operation that we exclude as biologically implausible. A more plausible method for extracting 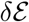 predates the Williams method (Barto et al., 1983; Doya and Sejnowski, 1988). It involves subtracting the average error from the trial-to-trial error. The residual of the subtraction is then simply the variation of the error 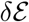 as desired.

As mechanism for this subtraction, we propose that the excitatory synaptic drive to the inferior olive is on average balanced by input from the GABAergic nucleo-olivary neurones. A diagram is shown in Fig. 8 to illustrate how this might work in the context of repetitions of a single movement (we extend the mechanism to multiple interleaved movements below). Briefly, a feedback plasticity reinforces the inhibition whenever it is too weak to prevent an error complex spike from being emitted. When the inhibition is strong enough to prevent an error complex spike, the inhibition is weakened. If the variations of the strength of the inhibition are sufficiently small, the level of inhibition provides a good approximation of the average error. Indeed, this mechanism can be viewed as maintaining an estimate of the movement error. However, the error still varies about its mean on a trial-to-trial basis because of the random perturbations that influence the movement and therefore the error. In consequence, error complex spikes are emitted when the error exceeds the (estimated) average; this occurs when the perturbation increases the error. This mechanism enables extraction of the sign of *δℰ* in the context of a single movement realisation. In support of such a mechanism, there is evidence that inhibition in the olive builds up during learning and reduces the probability of complex spikes (Kim et al., 1998).

**Figure 8:**
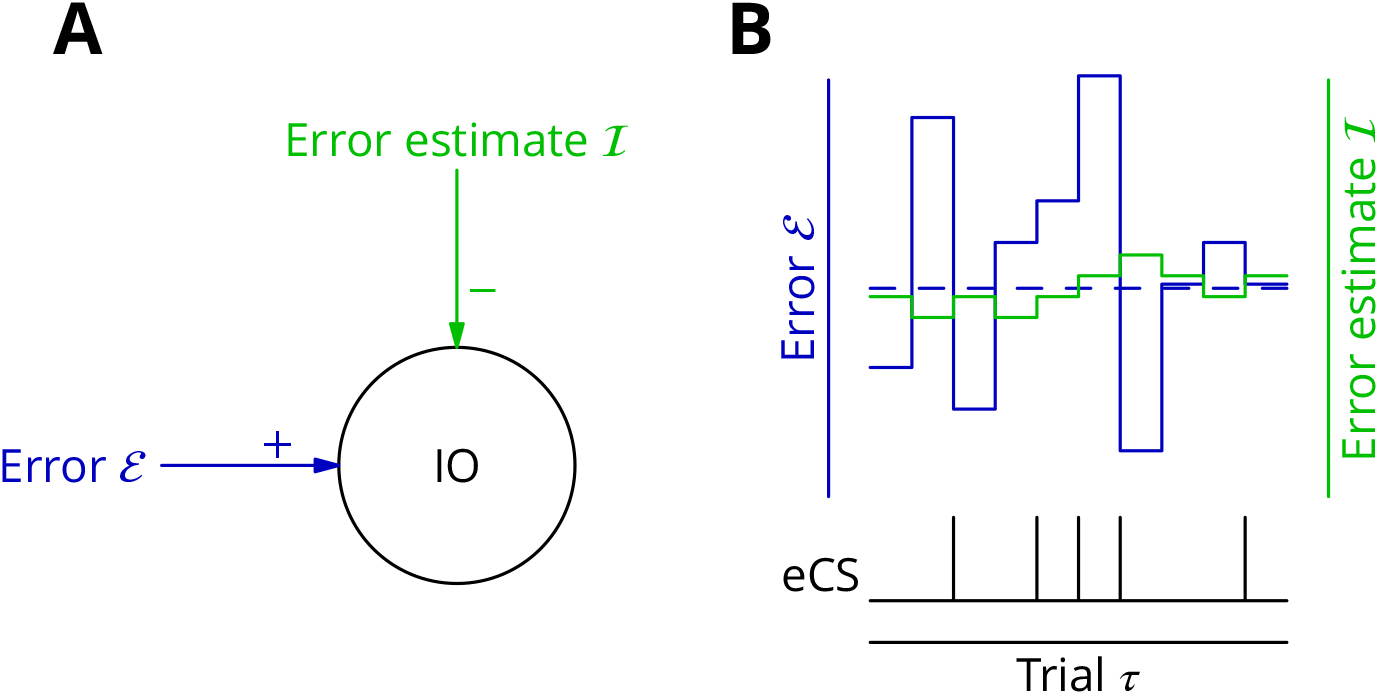
Adaptive tracking to cancel the mean error input to the inferior olive. **A**. The olive is assumed to receive an excitatory signal representing movement error 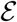 and an inhibitory input 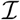 from the nucleo-olivary neurones of the cerebellar nuclei. **B**. The inputs to the inferior olive are represented in discrete time—each bar can be taken to represent a discrete movement realisation. The error (blue) varies about its average (dashed blue line) because perturbation complex spikes influence the movement and associated error randomly. The strength of the inhibition is shown by the green trace. When the excitatory error input exceeds the inhibition, an error complex spike is emitted (bottom black trace) and the inhibition is strengthened by plasticity, either directly or indirectly. In the converse situation and in the consequent absence of an error complex spike, the inhibition is weakened. In this way the inhibition tracks the average error and the emission of an error complex spike signals an error exceeding the estimated average. Note that spontaneous perturbation complex spikes are omitted from this diagram.

More than one plasticity mechanism could produce the desired cancellation of excitatory drive to the inferior olive. We outline two possibilities here, but it will be necessary in the implementation below to make a concrete if somewhat arbitrary choice; we shall make it on the basis of the available, circumstantial evidence.

The first possible mechanism would involve plasticity of the nucleo-olivary synapses. Perturbation and error complex spikes would be distinguished in an appropriate plasticity rule by the presence of excitatory synaptic input to the olive. This would offer a simple implementation, since plastic and cancelled inputs would be at neighbouring synapses (possibly even in the same olivary glomeruli: de Zeeuw et al., 1998); information about olivary spikes would also be directly available. However, the lack of published evidence and our own unsuccessful exploratory experiments led us to consider an alternative plasticity locus.

A second possible implementation for cancelling the average error signal would make the mossy fibre to nucleo-olivary neurone synapses plastic (Fig. 1B). The presence of an error complex spike would need to potentiate these inputs, thereby increasing inhibitory drive to the olive and tending to reduce the likelihood of future error complex spikes being emitted. Inversely, the absence of the error complex spike should depress the same synapses. Movement specificity could be conferred by applying the plasticity only to active mossy fibres, the patterns of which would differ between movements. This would enable movement-specific cancellation as long as the overlap between mossy fibre patterns was not too great.

How would information about the presence or absence of the error complex spike be supplied to the nucleo-olivary neurones? A direct connection between climbing fibre collaterals and nucleo-olivary neurones exists (de Zeeuw et al., 1997), but recordings of cerebellar neurones following stimulation of the olive suggests that this input is not strong, probably eliciting no more than a single spike per activation (Bengtsson et al., 2011). The function of this apparently weak input is unknown.

An alternative route to the cerebellar nuclear neurones for information about the error complex spike is via the Purkinje cells. Climbing fibres excite Purkinje cells which in turn inhibit cerebellar nuclear neurones, in which a strong inhibition can cause a distinctive rebound of firing (Llinás and Mühlethaler, 1988). It has been reported that peripheral stimulation of the climbing fibre receptive field, which might be expected to trigger the emission of error complex spikes, causes large IPSPs and an excitatory rebound in cerebellar nuclear neurones (Bengtsson et al., 2011). These synaptically induced climbing fibre–related inputs were stronger than spontaneously occurring IPSPs. In our conceptual framework, this could be interpreted as indicating that error complex spikes are stronger and/or arise in a greater number of olivary neurones than perturbation complex spikes. The two types of complex spike would therefore be distinguishable, at least in the cerebellar nuclei.

Plasticity of active mossy fibre inputs to cerebellar nuclear neurones has been reported which follows a rule similar to that our implementation requires. Thus, mossy fibres that burst before a hyperpolarisation (possibly the result of an error complex spike) that triggers a rebound have their inputs potentiated (Pugh and Raman, 2008), while mossy fibres that burst without a succeeding hyperpolarisation and rebound are depressed (Zhang and Linden, 2006). It should be noted, however, that this plasticity was studied at the input to projection neurones and not at that to the nucleo-olivary neurones. Nevertheless, the existence of highly suitable plasticity rules in a cell type closely related to the nucleo-olivary neurones encouraged us to choose the cerebellar nuclei as the site of the plasticity that leads to cancellation of the excitatory input to the olive.

We now consider how synaptic integration in the olive leads to emission or not of error complex spikes. The nucleo-olivary synapses (in most olivary nuclei) display a remarkable degree of delayed and long-lasting release (Best and Regehr, 2009), suggesting that inhibition would build up during a command and thus be able to oppose the excitatory inputs signalling movement errors that appear some time after the command is transmitted. The error complex spike would therefore be produced (or not) after the command. On this basis, we shall make the simplifying assumption that the cerebellum generates a relatively brief motor control output or ‘command’, of the order of 100 ms or less and a single error calculation is performed after the end of that command. As for the saccade example in the Introduction, many movements can only be evaluated after completion. In effect, this corresponds to an offline learning rule.

Climbing fibres are generally described as possessing highly specific modalities and receptive fields that are stereotypically linked to functional regionalisation in the cerebellum (Garwicz et al., 1998; Jörntell et al., 1996). Despite this, we shall make the simplifying assumption that all olivary cells serving a microzone transmit an error complex spike in a correlated manner. We return to the important issue of climbing fibre modalities in the Discussion.

## 7 Simulations

Above we outlined a mechanism for extracting the error change 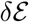 in the context of a single movement realisation; it is based on adapting the inhibitory input to the inferior olive to cancel the average excitatory error input in a movement-specific manner. To verify that this mechanism could operate successfully in conjunction with the scheme for cortical plasticity already described, we turned to simulation.

Before describing the model, it is important to make the point that any implementation will need to make concrete choices between alternative mechanisms that can in some cases be quite numerous and where the available evidence can only offer a poor guide, notably as concerns the properties of nuclear and olivary neurones and connections. Our aim here is therefore limited to showing that one specific implementation can implement stochastic gradient descent using cellular mechanisms that are plausible.

A reduced model of a cerebellar microzone was developed and is described in detail in §2.3. In overview, mossy fibre input patterns drove Purkinje and cerebellar nuclear neurones during commands of 10 discrete time bins. Purkinje cell activity was perturbed by randomly occurring complex spikes, which each increased the firing in a single time bin. The learning task was to adjust the output patterns of the nuclear projection neurones to randomly chosen targets. Cancellation of the average error was implemented by plasticity at the mossy fibre to nucleo-olivary neurone synapse while modifications of the mossy fibre pathway input to Purkinje cells reflected the rules outlined in §4. Synaptic weights were updated offline after each command. Error complex spikes were considered to be broadcast if the error exceeded the integral of inhibitory input to the olive during the movement. There were thus 400 (40 projection neurones × 10 time bins) variables to optimise using a single error value.

In a simplification that improved learning performance, the weight of the Purkinje cell–nucleo-olivary neurone connection was set to zero. This prevented the perturbations adding noise to the tracking of the average error. However, the connection was nevertheless assumed to be capable of signalling the presence of error complex spikes in order to guide the tracking plasticity. This apparent inconsistency could be resolved in vivo if the indirect effects of complex spikes were poorly transmitted to the inferior olive, possibly because of the specific dynamics of nucleo-olivary neurones and their output synapses (Najac and Raman, 2015; Best and Regehr, 2009). Moreover, our theoretical analysis below shows that constant inhibition of nucleo-olivary neurones is one mechanism allowing the maximum learning capacity of our algorithm to be attained.

The progress of the simulation is illustrated in Fig. 9, in which two different movements were successfully optimised in parallel; only one is shown. The global error is the sum of absolute differences between the projection neurone outputs and their target values. This can be seen to decrease throughout the simulation, indicating the progressive improvement of the learnt command. The effect of learning on the difference between the output and the target values can be seen in Fig. 9C and D. The initial match is very poor because of random initial weight assignments, but it has become almost exact by the end of the simulation.

**Figure 9:**
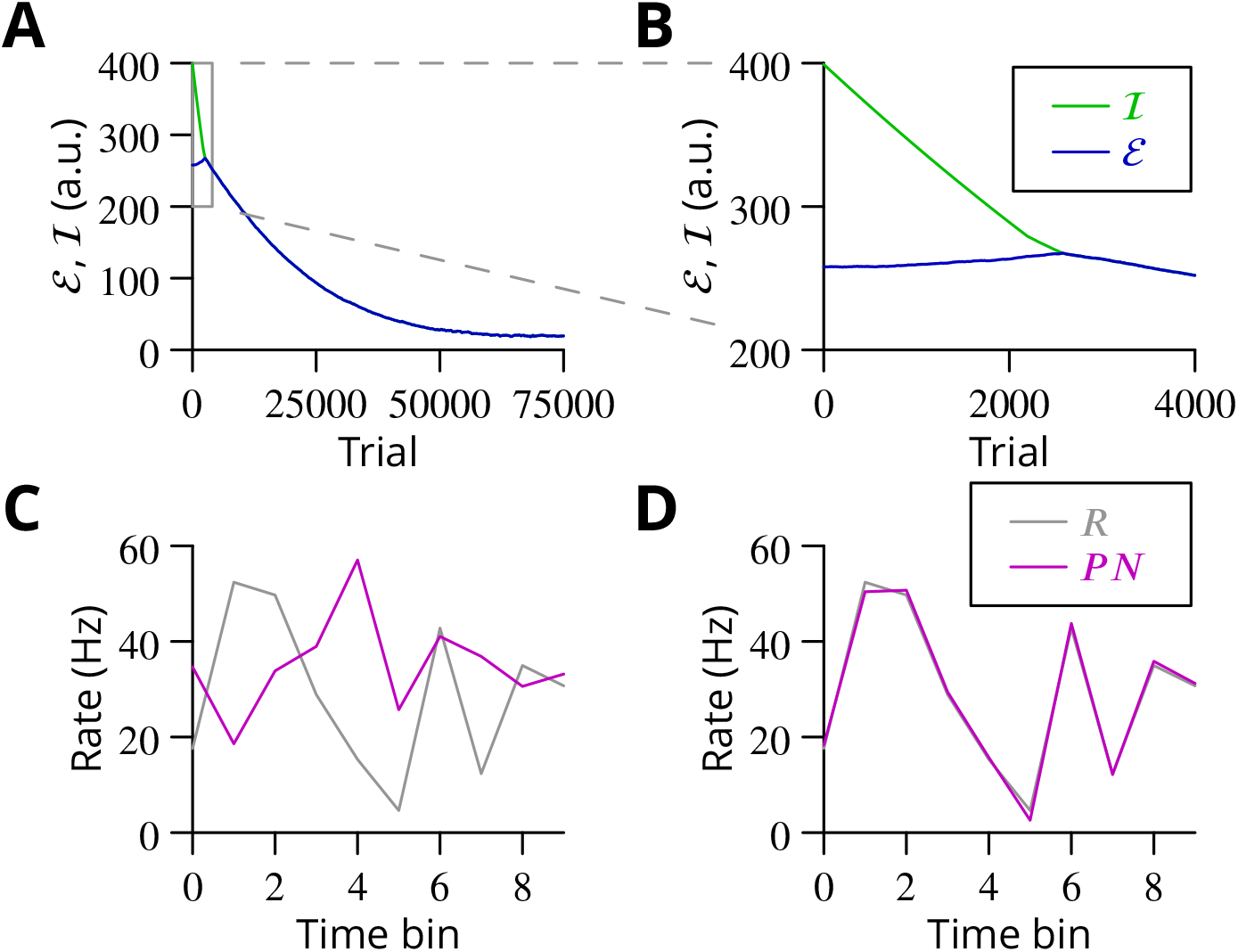
Simulated cerebellar learning by stochastic gradient descent with estimated global errors. The total error (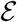, blue) at the cerebellar nuclear output and the cancelling inhibition (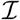, green) reaching the inferior olive are plotted as a function of trial number (*τ*) in **A** and **B** for one of two interleaved patterns learnt in parallel. An approximately 10-fold reduction of error was obtained. It can be seen in A that the cancelling inhibition follows the error very closely over most of the learning time course. However, the zoom in B shows that there is no systematic reduction in error until the inhibition accurately cancels the mean error. **C**. Initial firing profile of a typical cerebellar nuclear projection neurone (*PN*, magenta); the simulation represented 10 time bins with initially random frequency values per neurone, with a mean of 30 Hz. The target firing profile for the same neurone (*R*, grey) is also plotted. **D**. At the end of the simulation, the firing profile closely matched the target.

The optimisation only proceeds once the inhibitory input to the olive has accurately subtracted the average error. Thus, in Fig. 9B it can be seen that the initial values of the inhibitory and excitatory (error) inputs to the olive differed. The inhibition tends towards the error. Until the two match, the overall error shows no systematic improvement. This confirms the need for accurate subtraction of the mean error to enable extraction of the error changes necessary descend the error gradient. This simulation supports the feasibility of our proposed cerebellar implementation of stochastic gradient descent.

## 8 Stochastic gradient descent with estimated global errors

The simulations above provide a proof of concept for the proposed mechanism of cerebellar learning. Nonetheless, the specific scheme of stochastic gradient descent operating with estimated errors appears worthy of more detailed mathematical examination. Even the relatively simple network model in the simulations of §7 contains several parameters. It is by no means obvious to determine the regions of parameter space in which the model converges to the desired outputs or to find the parameter values that maximise convergence speed given a level of admissible residual error. To address these issues, we abstract the core mechanism of stochastic gradient descent with an estimated global error in order to highlight the role of four key parameters. Analysis of this mechanism will show that this algorithm, even in this very reduced form, exhibits a variety of dynamical regimes, which we characterise. We then show how the key parameters and the different dynamical learning regimes directly appear in an analog perceptron description of the type considered in the previous section, and show that the algorithm’s storage capacity is similar to the optimal capacity of an analog perceptron.

### 8.1 Minimal equivalent implementation

We begin by reducing the cerebellum-like scheme of Fig. 1B to its essence in Fig. 10. Purkinje cells and cerebellar nuclear projection neurones were combined into a single cell type, *P*. Learning a pattern (movement) requires adjustment of the rates *P_i_*(*t*), *i* = 1, ⋯, *N_P_* of *N_P_ P* cells to target values *R_i_, i* = 1, ⋯, *N_P_*. For now we consider only the rate of the *P* cell, without considering how it is produced (by synaptic inputs; the analysis will be extended below). The movement error 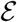 is defined as

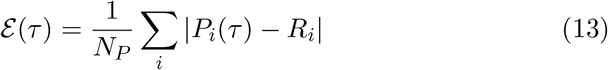

**Figure 10:**
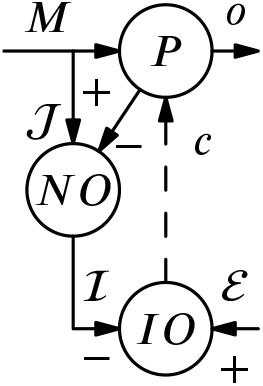
Simplified circuitry implementing stochastic gradient descent with estimated global errors. Purkinje cells and cerebellar nuclear projection neurones are collapsed into a single cell type *P* which provides the output *o*. These cells receive an excitatory plastic input from mossy fibres (*M*). Mossy fibres also drive a plastic inhibitory input 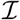, via the nucleo-olivary neurones, to the inferior olive (IO). The inferior olive receives an excitatory error input 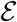. The olivary neurones emit spikes that are transmitted to the *P* cells via the climbing fibre *c*.

Note that here *τ* represents trial number rather than temporal variations within a single movement realisation. In this simplified model, *P* cell activity during a movement is characterised by a single value rather than a sequence of values in time.

The proposed core operation of the algorithm for one movement can be described as follows. At each time step, one cell *i_c_* is chosen at random and tested (perturbed by the climbing fibre): its firing rate is increased by *A* > 0 and becomes *P_i_c__* (*τ*) + *A*. The value of the global error corresponding to this perturbed firing rate is thus

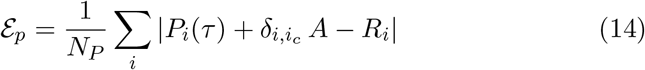

where we have used the Kronecker *δ_i,j_* notation (*δ_i,j_* = 1 if *i* = *j* and 0 otherwise).

The value of the perturbed error 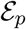 is compared to the current value of the estimated global error 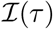, which is represented by the strength of the inhibitory *M* → *IO* input. Then *P_i_c__*(*τ*) and 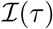 are updated depending on whether 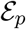 is smaller or greater than 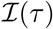.

- if 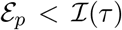 the perturbation is judged to have increased the error and therefore to have the wrong sign: the perturbed firing rate needs to be decreased. Concurrently, 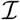 needs to be increased since it is judged too low compared to the real value of the error. Thus *P_i_c__*(*τ*) and 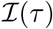 are changed to

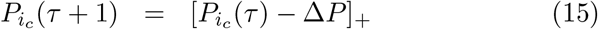

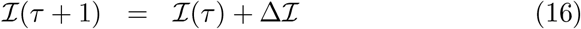

with the brackets indicating rectification (which imposes the constraint that firing rates are non-negative). The other rates are not changed *P_i_*(*τ* + 1) = *P_i_*(*τ*) for *i* ≠ *i_c_*
- if 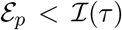, the converse reasoning leads to changes of *P_i_c__*(*τ*) and 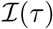 in opposite directions

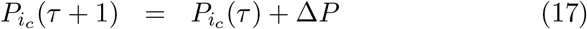

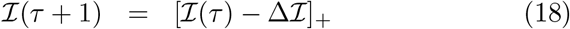

These updates by themselves are, however, not sufficient to lead to convergence. It is not difficult to verify that they preserve the quantity

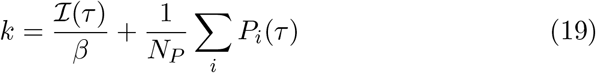

where we have introduced

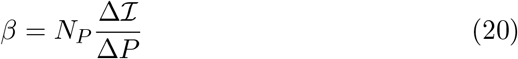

This conserved quantity prevents the learning dynamics from reaching the vicinity of the desired state 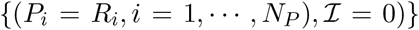 (when the value of k does not initially correspond to the specific desired final value). Minimal changes to the procedure are sufficient to remove this undesirable feature. One such modification that we here consider consists of performing the updates described by Equations (15-18), of ‘type 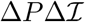’, only for a random fraction *ρ* of the time steps (*ρ* is the probability of a perturbation occurring in the trial). For the complementary (1 − *ρ*) fraction of time steps, updates of type 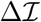 are performed as described above but without any perturbation (*A* = 0) and without any update of the *P_i_*s (only the estimated error 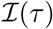 is updated in updates of type 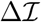).

This abstract model depends on the 4 parameters (besides the number of cells *N_P_*): *A*, the amplitude of the rate perturbation, Δ*P* and 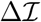, the update amplitudes of the rate and error estimate, respectively, and *ρ*, which describes the probability of a update of type 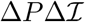 (and the complementary probability 1 − *ρ* of 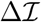 updates). The target firing rates are chosen in the range [0, *R_max_*], which simply fixes the firing rate scale. We would like to determine the conditions on the 4 parameters A, Δ*P*, 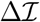 and *ρ* for the algorithm to ‘converge’^4^. We would also like to understand how they determine the rate of convergence and the residual error.

### 8.2 Convergence in the one-cell case

We analyse here the simplest case when the firing rate *P* of a single cell needs to be adjusted to reach a target value *R* (the case *N_P_* = 1 of the abstract model above). Stochastic gradient descent is conveniently analysed with a phase-plane description, by following the values 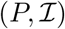 from one update to the next in the 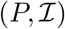 plane. The dynamics randomly alternates between updates of type 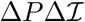 and 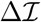, which we consider in turn.

For updates of type 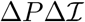, an update depends upon whether the perturbed error 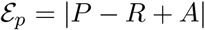 is larger or smaller than the estimated error 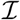. Namely, it depends on the location of the current 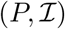 with respect to the two lines in the 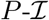 plane, 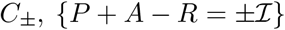, of slopes ±1 (see Figure 11). Note that the lines *D* define the error function. The dynamics of Equations (15-18) is such that each update moves the point 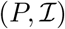 by adding to it the vectorial increment 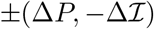 with the + sign holding in the quadrant above the lines *C*_±_ and the minus sign elsewhere. These updates move the point 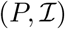 towards the lines *C*_±_.

**Figure 11:**
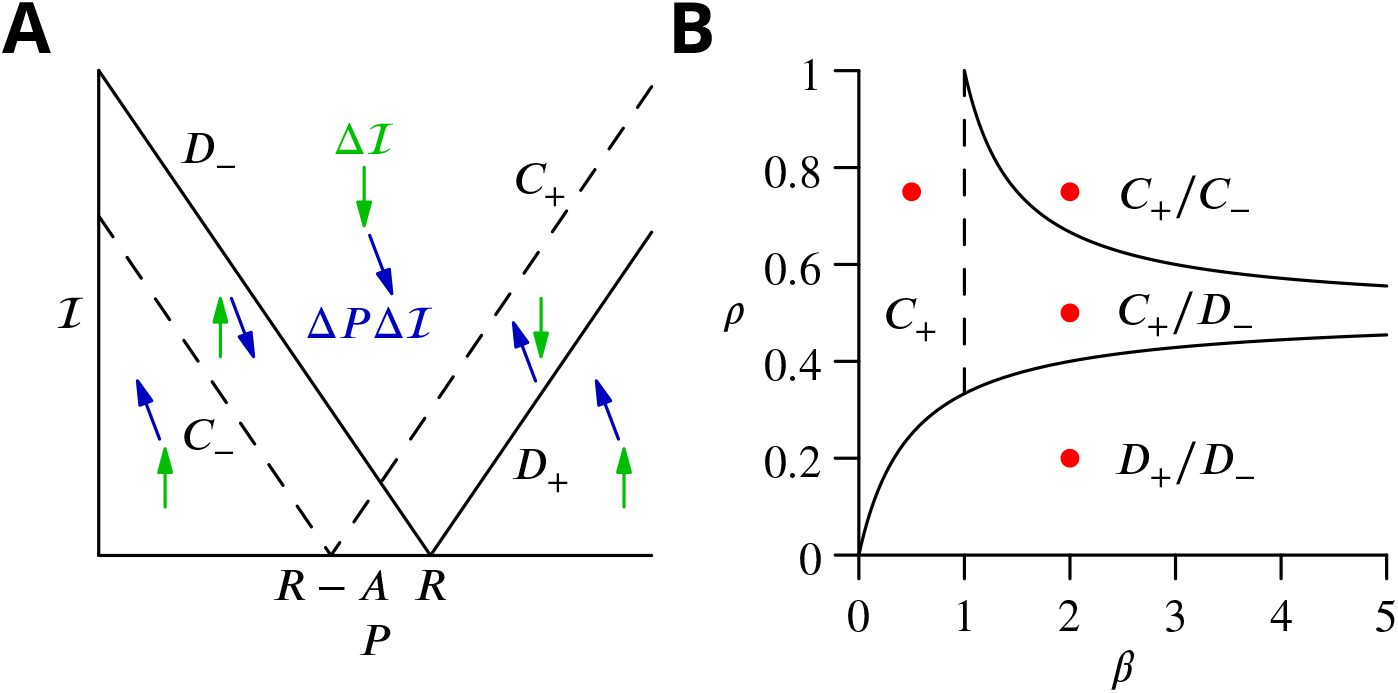
Reduced model: one-cell dynamics in the 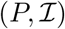 plane. **A**. The increment of updates of type 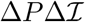 (blue arrows) and of type 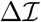 (green arrows) are shown. Updates of type 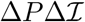 change sign when the *C*_+_ and *C*_−_ lines (dashed) are crossed. Updates of type 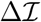 change sign when the *D*_+_ and *D*_−_ lines (solid) are crossed. The convergence of the rate *P* toward its target rate *R* (and the allied reduction of estimated error 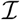) takes place as 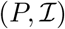 moves downward between the *C*_+_, *D*_+_ lines, i.e. the (*C*_+_, *D*_+_) corridor, or between the *C*_−_, *D*_−_ lines, i.e. the (*C*_−_, *D*_−_) corridor. **B**. When the perturbation amplitude is large enough, the convergence dynamics primarily takes place along one of the two lines of a convergence corridor. The chosen lines depend on the ratio 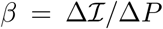 between the error and rate increment and on the probability *ρ* of choosing updates of type 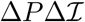. The diagram shows the selected lines depending on the values of *β* and *ρ*. The parameters of four cases illustrated in Fig. 12 are indicated (red dots).

Updates of type 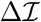 depend on the location of the point 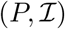 with respect to the lines *D*_±_, 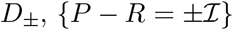, since the cell firing rate is not perturbed in these updates. In the quadrant above the lines *D*_±_, an update moves the point 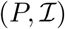 downwards by 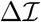, i.e. adds the vectorial increment 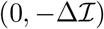. In the complementary domain of the 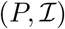 plane, an update moves the point 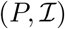 upwards by 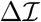, i.e. adds the opposite vectorial in-crement 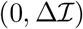. Both updates move 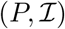 towards the lines *D*_±_.

Updates of type 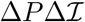 and 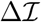 are performed with respective probabilities *ρ* and (1 − *ρ*). In the quadrant above the lines *C*_+_ and *D*_−_, these mixed updates lead the couple 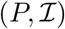 to perform a random walk with a systematic mean rightward-downward drift per update equal to

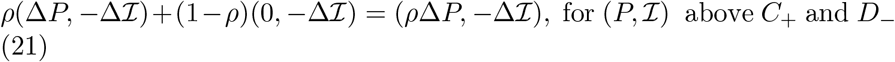

Below the lines *C*_−_ and *D*_+_, the updates are opposite and the mean drift per update is leftward-upward, equal to

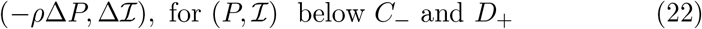

Depending on its initial condition and the exact set of drawn updates, this leads 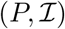 to reach either one of the two ‘convergence corridors’, between the lines *C*_+_ and *D*_+_, or between the lines *C*_−_ and *D*_−_ (see Figure 11)^5^.

Stochastic gradient descent dynamics thus proceeds in three successive phases, as seen in the simulations of §7. First, the firing rate and estimated error 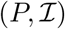 drift from the initial condition towards one convergence corridor. The duration of this phase depends on the location of the initial 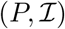 with respect to the convergence corridors and on the average drift (Equations (21) or (22)).

When a corridor is reached, in a second phase, 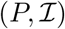 follows a stochastic walk in the corridor with, under suitable conditions, a mean linear decrease in time with bounded fluctuations. In a final phase, 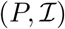 fluctuates around the intersection of *C*_+_ and *D*_−_, namely (*R* − *A*/2, *A*/2).

Convergence is thus controlled by the dynamics in the two corridors. A sufficient condition for convergence is that a single update cannot cross the two boundary lines of a corridor at once. The crossing of the (*C*_+_, *D*_+_) corridor by a single 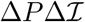 update provides the most stringent requirement,

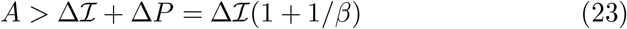

When this condition is met, alternation between the two types of updates produces a mean downward drift of the error. This downward drift controls the convergence rate and depends on the relative size of the perturbation and the discrete 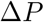 and 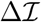 modifications (in other words, the number of modifications that are needed to cross the convergence corridor). For simplicity, we consider the case where the perturbation *A* is large compared to 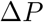 and 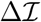, so that oscillations basically take place around one line of the corridor, as illustrated in Figure 12. This amounts to being able to neglect the probability of crossing the other line of the corridor. The convergence rate then depends on the corridor line around which it takes place and can be obtained by noting that around a given corridor line, one of the two types of updates always has the same sign.

**Figure 12:**
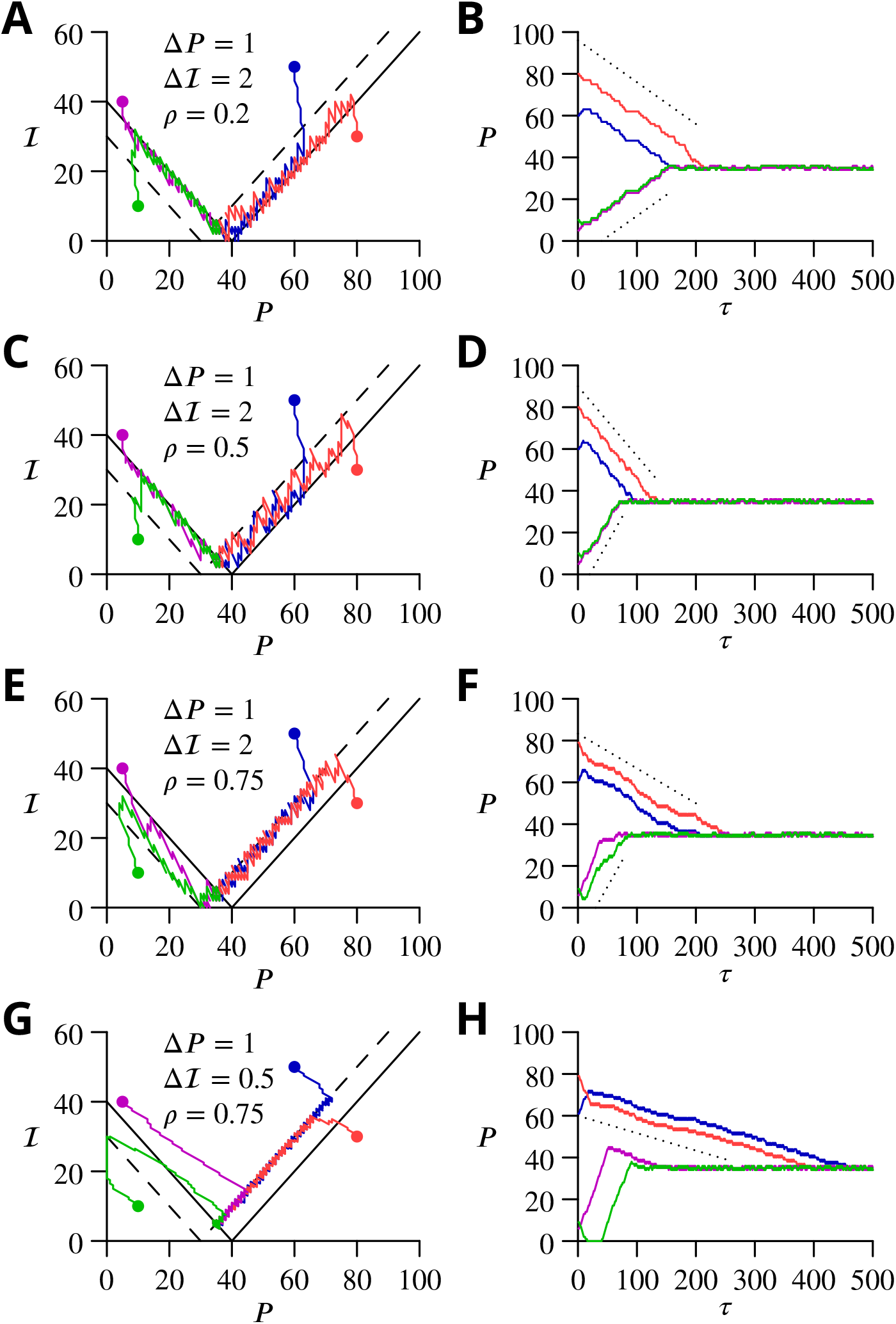
Reduced model: four cases of stochastic gradient descent learning for one cell. **A**. Dynamics in the 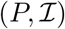 plane for *ρ* = 0.2 and 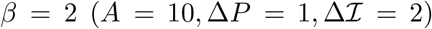. Trajectories (solid lines) from different initial conditions (filled circles) are represented in the 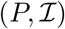 plane. The trajectories converge by oscillating around *D*_+_ in the *C*_+_/*D*_+_ ‘corridor’ and around *D*_−_ in the *C*_−_/*D*_−_ ‘corridor’. **B**. Time courses of *P*(*τ*) with corresponding colours. The slope of convergence (dotted lines) predicted by Equation (25, 26) agrees well with the observed convergence rate (although in the simulation the perturbation amplitude *A* is comparable to Δ*P* = 1 and 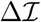). **C, D**. Same graphs for *ρ* = 0.5. The trajectories converge by oscillating around *C*_+_ in the *C*_+_/*D*_+_ ‘corridor’. **E, F**. Same graphs for *ρ* = 0.75. The trajectories converge by oscillating around *C*_−_ in the *C*_−_/*D*_−_ ‘corridor’. **G, H**. Same graphs for *ρ* = 0.75 and *β* = 0.5 (i.e. 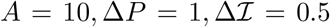). The trajectories converge by oscillating around *C*_+_, which is the only attractive line.

In the (*C*_+_, *D*_+_) corridor, performing type 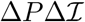 updates for fraction *ρ* of the steps and type 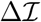 updates for the complementary fraction (1 − *ρ*) leads to the average displacement per step,

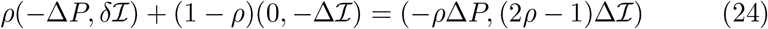

Thus, as summarised in Figure 11B

- for *ρ* < *β*/(1 + 2*β*) the average displacement leads to *D*_+_ (Figure 12A). The average displacement can be obtained by noting that for a large *A*, the 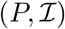 trajectory does not cross the *C*_+_ line^6^. Therefore, all the type 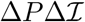 updates are of the same sign, of the form 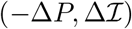, and chosen with probability *ρ*. Since updates of type 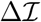 do not change the value of *P, P* approaches its target rate with a mean speed per step *V_c_*,

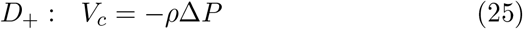 Comparison between this computed drift and simulated trajectories is shown in Figure 12B.
- for *ρ* > *β*/(1+2*β*), the average displacement leads to *C*_+_ (Fig. 12C,E,G). In this case, all type 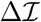 updates are of the same sign, of the form 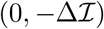. In contrast, a fraction *f* of type 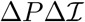 updates are of the form 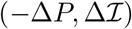, while a fraction (1 − *f*) is of the opposite form 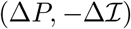 with *f* to be determined. The average drift is thus 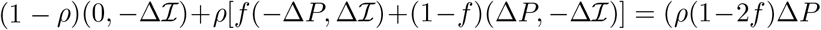, 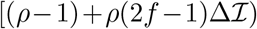. Requiring that this drift have slope 1, like *C*_+_, gives 2*f* − 1 = *β*(1 − *ρ*)/[(1 + *β*)*ρ*] and *f* = (*β* + *ρ*)/[2*ρ*(1 + *β*)] (which indeed obeys 0 < *f* < 1 in the parameter domain considered). Thus, *P* approaches its target rate with a mean speed per step *V_c_*,

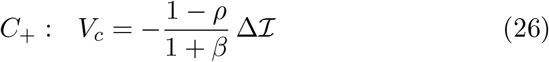 Comparison between this computed drift and simulated trajectories is shown in Fig. 12D,F,H.

In the (*C*_−_, *D*_−_) corridor, the average drift per step is opposite to the drift in the (*C*_+_, *D*_+_) corridor (Equation (24)). Thus,

- for *β* > 1 and *ρ* > *β*/(2*β* − 1) the average displacement leads to *C*_−_ (Fig. 12E). Updates of type 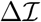 are always of the form 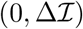, while a fraction *f* of type 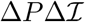 updates are of the form 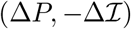 and a fraction (1 − *f*) are of the opposite form 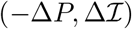. Again, since the average drift is along *C*_−_ of slope −1, one obtains 2*f* − 1 = *β*(1 − *ρ*)/*β* − 1)*ρ*] or *f* = (*β* − *ρ*)/[2*ρ*(*β* − 1) (which obeys 0 < *f* < 1 in the parameter domain considered). The convergence speed *V_c_* is thus,

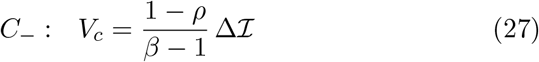 Comparison between this computed drift and simulated trajectories is shown in Fig. 12F.
- for *β* < 1, or (*β* > 1, *ρ* < *β*/(2*β* − 1)), the complementary parameter domain, the drift in the corridor leads to *D*_−_ (Fig. 12A,C). The domain *β* < 1 can be excluded since when the point 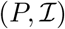 crosses *D*_−_, the drift in the upper quadrant (*C*_+_, *D*_−_) tends to bring it to the other corridor (*C*_+_, *D*_+_)(Fig. 12G). Near the line *D*_−_, type 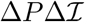 updates are always of the form 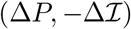. The convergence speed *V_c_* is thus

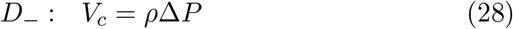

Comparison between this computed drift and simulated trajectories is shown in Fig. 12B,D.

The expressions obtained for the convergence rates allow us to address the question of parameter optimisation and the fastest convergence rate for a final given error (assumed for now to be determined by *A* namely an error of about *A*/2).

We consider first the (*C*_+_, *D*_+_) corridor and optimise the three parameters *ρ*, 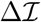 and Δ*P* in turn.

Since *V_c_* is linear in *ρ* for convergence around the two lines, the maximal |*V_c_*| is obtained by maximising *ρ* for convergence around *D*_+_ or maximising it for convergence around *C*_+_. In both cases, the maximal speed is obtained at the boundary of the respective parameter domains *ρ* = *β*/(1 + *β*) and is equal to

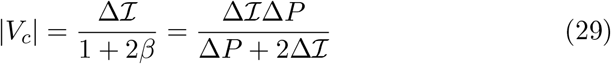

We would like to optimise 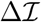 and Δ*P* taking into account the convergence constraint 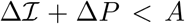. Unfortunately, we cannot rigorously do this, since we have determined the convergence rate only for 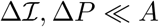, Δ*P* ≪ *A*. Therefore, we shall assume that the expression (29) is approximately correct in the whole convergence domain and make use of it to determine the optimal values of 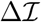 and Δ*P*. Since |*V_c_*| in Equation (29) is homogeneous of degree 1 in 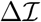 and Δ*P*, its maximal value is obtained on the convergence domain boundary 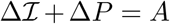. Then, an elementary maximisation yields the optimal parameters

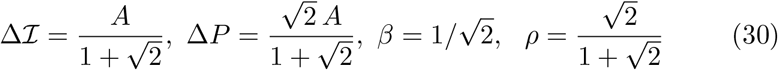

and the maximal convergence rate in the (*C*_+_, *D*_+_) corridor

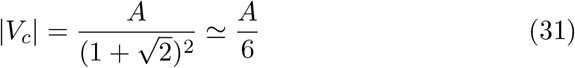

For these parameters, since *β* < 1, the (*C*_−_, *D*_−_) corridor is not a stable attractor and need not be considered.

A similar optimisation for the (*C*_−_, *D*_−_) corridor alone would give as limiting optimal values 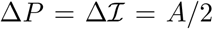, together with *ρ* = 1 and a rate |*v_c_*| = *A*/2. However, these parameters give a vanishingly slow convergence along the *C*_+_ line, in the other corridor. Thus, the choice of optimal parameters will depend on the statistics of the initial condition 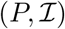 and how different rates are combined, but choices intermediate between these values and (30) are likely to yield the fastest overall convergence rate under most conditions.

### 8.3 Convergence and capacity in an analog perceptron model

In order to show how the previous results map onto a model of the type simulated in §7, we consider learning of multiple associations for an analog perceptron with the algorithm of stochastic gradient descent with estimated global errors.

The architecture is again the simplified circuit of Fig. 10, with *N_M_* = 1000 mossy fibres projecting onto a single *P* cell, with weights *w_i_, i* = 1, …, *N_M_* (which are all positive or zero). *N_P_* random patterns are generated randomly. Activities of mossy fibres in different patterns are i.i.d. binary random variables 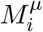 with coding level *f* (i.e. *M_i_* = 1 with probability *f* and 0 with probability 1 − *f*; in the present simulations *f* = 0.2). The Purkinje cell desired rates for the *N_P_* patterns are i.i.d. uniform variables *R^μ^* from 0 to *P_max_*.

In the case of the stochastic gradient descent rule, at each trial there is a probability *ρ* of a perturbation of amplitude *A*, so that

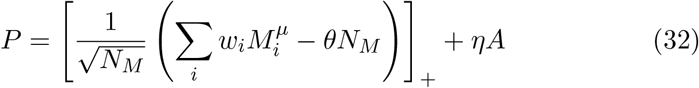

where *η* = 1 with probability *ρ* and *η* = 0 otherwise, which thus introduces a random perturbation of amplitude *A* into the *P*-cell firing. The error is defined as 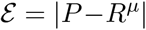. Comparison with previously obtained results for the capacity of this analog perceptron (Clopath and Brunel, 2013) motivates our choice of weights normalisation and the parameterisation of the threshold as 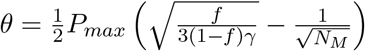, where *γ* is a composite parameter reflecting the statistics of input and output firing, but here is equal to one.

An inferior olivary neuron receives the excitatory error signal but it also receives inhibitory inputs from the nucleo-olivary neurones driven by the mossy fibre inputs (which we have denoted *M* above), with weights *v_i_*. These are also plastic and sign-constrained. They represent the current estimated error. The net drive of the inferior olivary neurone is

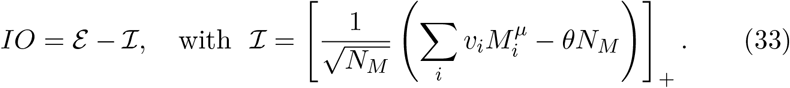

Here subtracting *θ* represents the inhibition of nucleo-olivary neurones by Purkinje cells. For simplicity we assume this inhibition is constant and the value of *θ* is equal to that used in Equation (32).

The climbing fibre signal controlling plasticity is

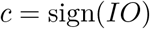

Weights are changed according to the following rule. Weights of *M* → *P* synapses active simultaneously with a perturbation are increased if *c* is negative and decreased if it is positive. Weights of *M* → *IO* synapses are increased if *c* is positive and decreased if not. Thus

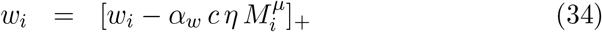

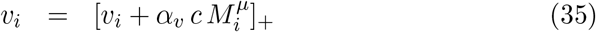

with the brackets indicating rectification (to impose the excitatory constraint).

The parameters of the simplified model of the previous subsection can be written as a function of those controlling the learning process in the present analog perceptron. The probability *ρ* of the two types of updates (with and without perturbation) and the amplitude of perturbation *A* are clearly identical in both models. In order to relate the previous Δ*P* and 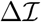 to the present amplitude change of the weights *α_w_* at mossy fibre–*P* cell inputs and *α_v_* for the indirect drive to the inferior olive from mossy fibres, we note that far below maximal learning capacity, the number of silent synapses is presumably small and the rectification constraints can be neglected in Equations (34,35). Therefore, the weight modifications result in the changes Δ*P*, of the perceptron firing rate, and 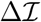, of the inhibitory input 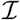 to the olive,

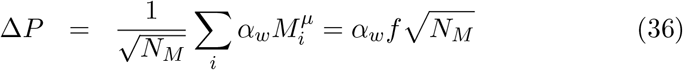

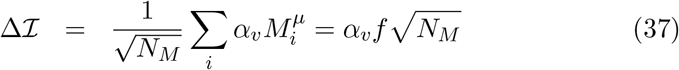

The convergence rate estimates of the previous section are compared with direct stochastic gradient descent learning simulations of the analog perceptrons in Fig. 13. As shown in Fig. 13A, the convergence rate for single patterns agrees well with the estimate (26)^7^. For a larger number of patterns below the maximal learning capacity, the convergence rate per pattern is slower by a factor of ≈ 1.5 − 5 for *N_P_* between 100 − 350 (Fig. 13A,C). This convergence is considerably slower than that obtained for the perceptron learning rule (Fig. 13B), but note that the relative slowing down due to interference of multiple patterns is comparable for the perceptron and the proposed algorithm (compare panels B,C in Fig. 13).

**Figure 13:**
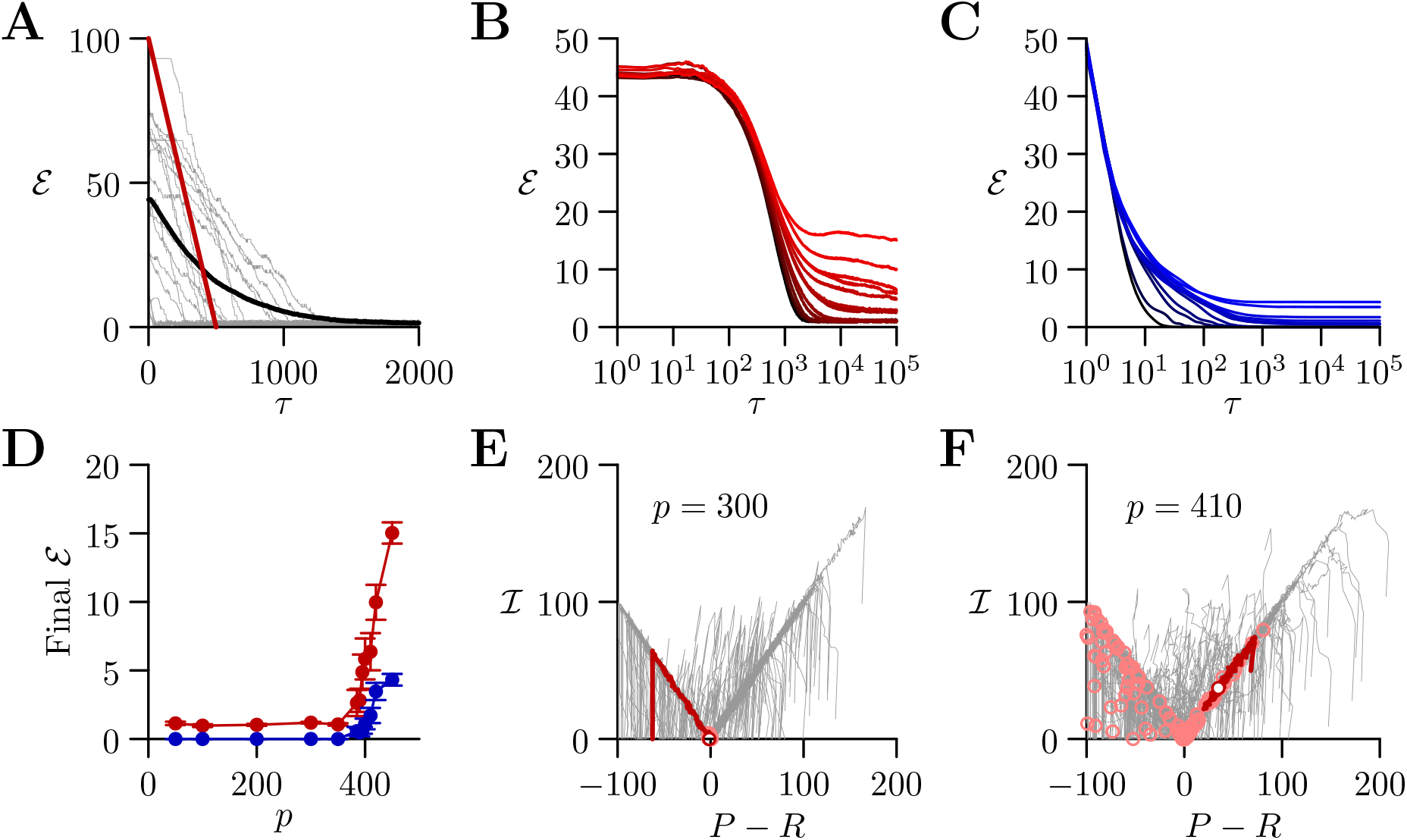
Convergence and capacity for an analog perceptron. **A**. Convergence for a single learned pattern (error 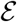 vs. number of trials, *τ*) for stochastic gradient descent with an estimated global error (SGDEGE). Grey: examples of individual realisations. Black: average error (*n* = 100). Red: predicted rate of convergence (see text §8.3) **B**. Mean error vs. number of trials (always per pattern) for different numbers of patterns for SGDEGE. Colours from black to red: 50, 100, 200, 300, 350, 385, 390, 395, 400, 410, 420, 450 patterns. **C**. Mean error vs. number of trials when learning using the perceptron rule. The final error is zero when the number of patterns is below capacity (predicted to be 391 for these parameters), up to finite size effects. Colours from black to blue: same numbers of patterns as listed for B. **D**. Mean error of the perceptron rule (blue, after 10^5^ trials) and the SGDEGE (red, after 10^5^ trials), as a function of the number of patterns *p*. The mean error diverges from its minimal value close to the theoretical capacity for both learning algorithms. **E, F**. Dynamics of pattern learning in the 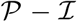 plane (*P* is the *P*-cell rate and *R* its optimum; 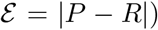 below (E) and above (F) capacity. Below capacity the error corresponding to each pattern (grey) is reduced as it moves down its convergence corridor. Above capacity, patterns cannot all converge. The final coordinates in the simulation are indicated with pink open circles (in E all are within *A* of (0,0)). In each of panels E and F, a specimen learning trajectory is shown in red with an open circle end-point.

Furthermore, we found that including the effect of *P*-cell inhibition of the nucleo-olivary neurones increases the capacity of the proposed algorithm by reducing the interference of error estimates corresponding to different patterns. The final average error increases from the baseline established for a single pattern (*A*/2 = 1) only very close to the theoretical capacity (Fig. 13D) which was computed analytically given the input-output association statistics (Clopath and Brunel, 2013).

Looking at the simultaneous convergence of patterns in the 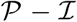 plane (Fig. 13E,F) we see that above capacity the interference between learning different patterns causes some of them to get ‘stuck’ with large errors.

In summary, SGDEGE converges to a finite error *A*/2 rather than zero, is slower than the perceptron algorithm, which exploits biologically-unrealistic complete error information, but can approach very closely the maximal storage capacity.

## 9 Discussion

### 9.1 A cellular implementation of stochastic gradient descent

Analysis of the requirements and constraints for a general cerebellar learning algorithm highlighted the fact that the current consensus Marr-Albus-Ito model is only capable of learning simple reflex movements. Optimisation of complex, arbitrary movements, of which organisms are certainly capable and to which the cerebellum is widely believed to contribute, would require a different algorithm. We therefore sought to identify within the cerebellar system an implementation of stochastic gradient descent. This should comprise several elements: a source of perturbations, a mechanism for extracting the change of error, and a plasticity rule incorporating this information. We identified a strong constraint on any implementation requiring each calculation to be made in the context of a single movement realisation. This arises from the potentially arbitrary sequencing of movements with different optima. We also sought a mechanism that only makes use of plausible cellular calculations: summation of excitation and inhibition in the presence of a threshold.

We suggest that the perturbation is provided by the complex spike, which has suitable properties: spontaneous irregular activity, an unambiguous sign, salience at a cellular and network level, and the ability to influence synaptic plasticity. This choice of perturbation largely determines the predicted cerebellar cortical plasticity rules: only granule cell inputs active at the same time as a perturbation complex spike undergo plasticity, whose sign is determined by the absence (LTP) or presence (LTD) of a succeeding error complex spike. We have verified that the synaptic plasticity rules do operate as predicted, in vitro under conditions designed to be more physiological than is customary.

A more involved mechanism seems to be required to read off the change of error. The general mechanism we propose involves subtraction of the average error to expose the random variations caused by the perturbations of the movement. The subtraction results from adaptive tracking of the excitatory input to the olivary neurones by the inhibitory input from the nucleo-olivary neurones of the cerebellar nuclei. We chose to place the plasticity at the mossy fibre–nucleo-olivary neurone synapse, mostly because of the existence of suitable plasticity rules at the mossy fibre synapse onto the neighbouring projection neurones. However, plasticity in the olive at the nucleo-olivary input would probably be functionally equivalent and we do not intend to rule out this or alternative sites of the error-cancelling plasticity.

By simulating a simplified cerebellar network implementing this mechanism, we established the ability of our proposed mechanism to learn multiple arbitrary outputs, optimising 400 variables per movement with a single error value. More formal analysis of a simplified version of stochastic gradient descent with estimated global errors established convergence of the algorithm and allowed us to estimate its storage capacity. The analysis also highlighted a potential benefit of retaining a low probability of complex spike emission—to enable the tracking plasticity to occur independently of the plasticity at parallel fibre–Purkinje cell synapses.

### 9.2 Implications for studies of synaptic plasticity

The plasticity rules for parallel fibre–Purkinje cell synapses predicted by our algorithm appeared to be almost completely incompatible with the well established consensus. However, once we eliminated known deviations from physiological conditions, we were able to verify the four predicted outcomes.

We made several changes to the experimental conditions, only one of which is specific to the cerebellum. One—leaving synaptic inhibition intact—has long been recognised as being of potential importance, with debates regarding its effect in hippocampal LTP dating back decades (Wigström and Gustafsson, 1983a,b; Arima-Yoshida et al., 2011).

We also made use of a lower extracellular calcium concentration than those almost universally employed in studies of plasticity in vitro. In vivo measurements of the extracellular calcium concentration suggest that it does not exceed 1.5 mM in rodents, yet most studies use at least 2 mM. A 25% alteration of calcium concentration could plausibly change plasticity outcomes, given the numerous nonlinear calcium-dependent processes involved in synaptic transmission and plasticity (Nevian and Sakmann, 2006; Graupner and Brunel, 2007).

A major change of conditions we effected was cerebellum-specific. Nearly all studies of granule cell–Purkinje cell plasticity have employed stimulation of parallel fibres in the molecular layer. Such concentrated, synchronised input activity is unlikely ever to arise physiologically. Instead of this, we stimulated in the granule cell layer, a procedure expected to generate a much more spatially dispersed input on the Purkinje cell, presumably leading to minimised dendritic depolarisations. Changing the stimulation method has been reported to prevent induction of LTD using standard protocols (Marcaggi and Attwell, 2007).

Although we cannot predict in detail the mechanistic alterations resulting from these changes of conditions, it is nevertheless likely that intracellular calcium concentrations during induction will be reduced, and most of the changes we observed can be interpreted in this light. It has long been suggested that high calcium concentrations during induction lead to LTD, while lower calcium concentrations generate LTP (Coesmans et al., 2004); we have recently modelled the induction of this plasticity, incorporating both calcium and nitric oxide signalling (Bouvier et al., 2016). Consistently with this viewpoint, protocols that under standard conditions produce LTD—simultaneous activation of granule cells and climbing fibres—could plausibly produce LTP in the present conditions as a result of reduced intracellular calcium. Analogously, granule cell stimulation that alone produces LTP under standard conditions might elicit no change if calcium signalling were attenuated under our conditions.

Interestingly, LTP resulting from conjunctive granule cell and climbing fibre stimulation has been previously reported, in vitro (Mathy et al., 2009; Suvrathan et al., 2016) and in vivo (Wetmore et al., 2014). In contrast, our results do not fit well with several other studies of plasticity in vivo (Ito et al., 1982; Jörntell and Ekerot, 2002, 2003, 2011). However, in all of these studies quite intense stimulation of parallel and/or climbing fibre inputs was used, which may result in greater depolarisations and calcium entry than usually encountered. This difference could therefore account for the apparent discrepancy with the results we predict and have found in vitro.

In summary, while in vitro studies of plasticity under unphysiological conditions are likely to reveal the molecular mechanisms leading to potentiation and depression, the precise outcomes from given stimulation protocols may be difficult to extrapolate to the in vivo setting, as we have shown here for the cerebellum. Similar arguments could apply to in vitro plasticity studies in other brain regions.

### 9.3 Current evidence regarding stochastic gradient descent

As mentioned in the introductory sections, the general cerebellar learning algorithm we propose here is not necessarily required in situations where movements are simple or constrained, admitting a fixed mapping between errors and corrective action. Furthermore, such movements constitute the near totality of well studied models of cerebellar learning. Thus, the vestibulo-ocular reflex and saccade adaptation involve eye movements, which are naturally constrained, while the eyeblink is a stereotyped protective reflex. There is therefore a possibility that our mechanism does not operate in the cerebellar regions involved in ocular motor behaviour even if it does operate elsewhere.

In addition, these ocular behaviours apparently display error functions that are incompatible with our assumptions (see Fig. 3C). In particular, disturbance of a well optimised movement would be expected to increase error. However, it has been reported multiple times that climbing fibre activity can provide directional error information, including *reductions* of climbing fibre activity below baseline (e.g. Soetedjo et al., 2008); this would be reminiscent of the curves of Fig. 3A. This argument is not totally conclusive, however.

Firstly, we recall that the error is represented by the input to the inferior olive, not its output. It is thus possible that inputs from the nucleo-olivary neurones (or external inhibitory inputs) to the olive also have their activity modified by the disturbance of the movement, causing the reduction of climbing fibre activity. Secondly, what matters for our algorithm is the temporal sequence of perturbation and error complex spikes, but these second-order statistics of complex spike activity have never been investigated. Similarly, it has been reported that learning and plasticity (LTD) occur in the absence of *modulation* of climbing fibre activity (Ke et al., 2009). Although this is difficult to reconcile with either the standard theory or our algorithm, it does not entirely rule out the existence of perturbation-error complex spike pairs that we predict lead to LTD.

Beyond the predictions for the plasticity rules at parallel fibre–Purkinje cell synapses tested above, there are a number of aspects of our theory that do fit well with existing observations. The simple existence of spontaneous climbing fibre activity is one. Additional suggestive features concern the evolution of climbing fibre activity during eyeblink conditioning (Ohmae and Medina, 2015). Once conditioning has commenced, the probability of complex spikes in response to the unconditioned stimulus decreases, which would be consistent with the build up of the inhibition cancelling the average error signal in the olive. Furthermore, omission of the unconditioned stimulus then causes a reduction in the probability of complex spikes below the baseline rate, strongly suggesting a specifically timed inhibitory signal has indeed developed at the time of the unconditioned stimulus (Kim et al., 1998).

We suggest that the cancellation of average error involves plasticity at mossy fibre–nucleo-olivary neurone synapses. To date no study has reported such plasticity, but the nucleo-olivary neurones have only rarely been studied. Plasticity at the mossy fibre synapses on projection neurones has been studied both in vitro (Pugh and Raman, 2006, 2008; Zhang and Linden, 2006) and in vivo (Ohyama et al., 2006), but is not used in our proposed algorithm. Axonal remodelling and synaptogenesis of mossy fibres in the cerebellar nuclei may underlie this plasticity (Kleim et al., 2002; Boele et al., 2013; Lee et al., 2015) and could also contribute to the putative plasticity at mossy fibre synapses on nucleo-olivary neurones.

Finally, our theory of course predicts that perturbation complex spikes perturb ongoing movements. It is well established that climbing fibre activation can elicit movements (Barmack and Hess, 1980; Kimpo et al., 2014; Zucca et al., 2016), but it remains to be determined whether the movements triggered by spontaneous climbing fibre activity are perceptible. Stone and Lisberger (1986) reported the absence of complex-spike-triggered eye movements in the context of the vestibulo-ocular reflex. However, it is known that the visual system is very sensitive to retinal slip (Murakami, 2004), so it may be necessary to carry out high-resolution measurements and careful averaging to confirm or exclude the existence of perceptible movement perturbations.

### 9.4 Climbing fibre receptive fields and the bicycle problem

There is an extensive literature characterising the modalities and receptive fields of climbing fibres. The great majority of reports are consistent with a view according to which climbing fibres have fixed, specific modalities or very restricted receptive fields, with neighbouring fibres having similar properties (Garwicz et al., 1998; Jörntell et al., 1996). Examples would be a climbing fibre driven by retinal slip in a specific direction (Graf et al., 1988) or responding only to a small patch of skin (Garwicz et al., 2002). These receptive fields are quite stereotyped and have proven to be reliable landmarks in the functional regionalisation of the cerebellum; they are moreover tightly associated with the genetically specified zebrin patterning of the cerebellum (Schonewille et al., 2006b; Mostofi et al., 2010; Apps and Hawkes, 2009).

The apparently extreme specialisation of climbing fibres implies a hitherto unrecognised limitation of the learning abilities of the cerebellum. This arises from the inescapable fact that a portion of the cerebellum receiving a single, specific type of error information can only optimise a movement with the aid of that feedback and none other. Thus, a cerebellar region controlling an animal’s forelimb might typically receive climbing fibres driven by cutaneous and proprioceptive input from part of that limb (Garwicz et al., 2002). This would enable learning of simple withdrawal reflexes, but would be unable to contribute to the learning of tasks guided by other error signals.

We can illustrate this with a more human behaviour: riding a bicycle, which is often taken as an example of a typical cerebellar behaviour. This is an acquired skill for which there is little apparent evolutionary precedent. It is likely to involve learning somewhat arbitrary arm movements in response to vestibular input (it is possible to ride a bike with one’s eyes closed). The error signals guiding learning could be vestibular, visual or possibly cutaneous/nociceptive (as a result of a fall), but not necessarily those related to the arm whose movement is learnt. How can such disparate or uncommon but sometimes essential error signals contribute to cerebellar control of the arm? We call this the ‘bicycle problem’.

At least two, non-exclusive solutions to this problem can be envisaged. The first we term ‘output convergence’; it involves the convergence of multiple cerebellar regions receiving climbing fibres of different modalities onto each specific motor element (for instance a muscle) being controlled. Striking, if partial, evidence for this is found in a study by Ruigrok et al. (2008), who injected the retrograde, trans-synaptic tracer rabies virus into individual muscles. Multiple cerebellar zones were labelled, showing that they all contribute to the control of those muscles, as posited. What is currently less clear is whether such separate zones receive climbing fibre inputs with different modalities. We note that the output convergence solution to the bicycle problem implies that the synaptic changes in those regions receiving the appropriate error information must outweigh any drift in synaptic weights from those regions deprived of meaningful error information. This could be partially implemented by adaptation of the learning rate, an algorithmic extension we suggest below.

**Figure 14:**
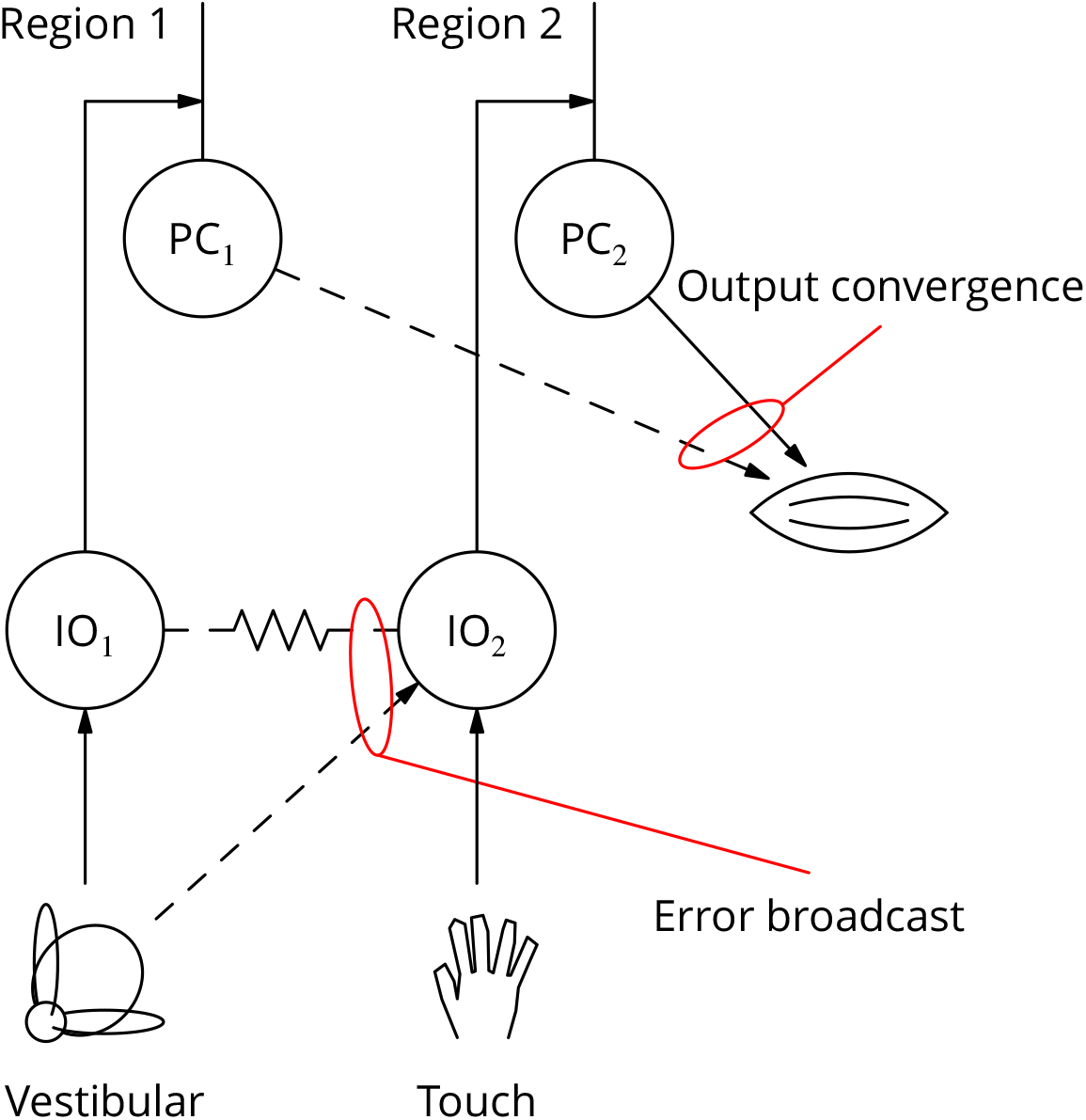
Diagram illustrating two possible solutions to the ‘bicycle problem’: how to use vestibular error information to guide learning of arm movements to ride a bicycle. In the ‘output convergence’ solution, the outputs from cerebellar regions receiving different climbing fibre modalities converge onto a motor unit (represented by a muscle in the diagram). In the ‘error broadcast’ solution, error complex spikes are transmitted beyond their traditional receptive fields, either by divergent synaptic inputs and/or via the strong electrical coupling between inferior olivary neurones.

We term the second solution to the bicycle problem the ‘error broadcast solution. According to this, error inputs to the olive are broadcast to olivary neurones (and Purkinje cells) outside the traditional receptive field. Although the weight of literature appears to be against this, there are both possible mechanisms and a small amount of supporting data for this suggestion. In terms of mechanism, divergence of synaptic inputs is of course a possibility, but there is additionally the well known electrical coupling of olivary neurones (Devor and Yarom, 2002). Both of these mechanisms could contribute to the recruitment of cells outside the traditional receptive fields. This may occur much more frequently in vivo than in the quiescent/anæsthetised conditions employed for most studies of climbing fibre receptive fields. Evidence for ‘broadcast’ of what we would term error complex spikes in vivo involves auditory and visual stimuli (Mortimer, 1975; Ozden et al., 2012); these stimuli may be correlated with startle responses. Eyeblink conditioning using a visual unconditioned stimulus has also been reported (Rogers et al., 1999).

The existence of broadcast error complex spikes would provide a mechanism explaining the giant IPSPs in the cerebellar nuclei elicited by peripheral stimulation (Bengtsson et al., 2011) and could also account for the correlation between visual smooth pursuit behaviour and the nature of individual complex spikes (Yang and Lisberger, 2014): behaviour could only correlate with a single cell if others were receiving the same input.

### 9.5 Possible extensions to the algorithm

Our implementation of cerebellar stochastic gradient descent and its simulation were purposefully kept as simple as possible, to provide a proof-of-concept with a minimum of assumptions and to simplify parallel analyses. It is likely that parts of the implementation will need to be altered and/or extended as further information becomes available.

Probably the most uncertain element of the implementation is the adaptive site in the cancellation of the average error. We chose to make the mossy fibre–nucleo-olivary neurone synapse plastic, but the plasticity could certainly operate in the olive instead of or in addition to the cerebellar nuclear site. Further studies of synaptic transmission and plasticity in both structures are clearly warranted in this context.

A simplification in our implementation is that it represents brief, discrete commands in what amounts to an offline learning rule. Error complex spikes are only emitted after the command and indeed were not simulated explicitly. This has the great advantage of avoiding the question of whether Purkinje cells that have not received a perturbation complex spike would interpret a broadcast error complex spike as a perturbation. We believe that plausible cellular mechanisms exist that would enable the Purkinje cell to distinguish the two types of input. The most obvious would be that, as already hinted at in the literature, error complex spikes are likely to be stronger. An extended synaptic plasticity rule could therefore include a case in which an error complex spike received in the absence of a recent perturbation spike has a neutral plasticity effect. There is currently little data on which to base a detailed implementation.

An open question is whether different relative timings of parallel and climbing fibre activity would result in different plasticity outcomes. In particular, one might hypothesise that parallel fibres active during a pause following a perturbation complex spike might display plasticity of the opposite sign to that reported here for synchrony with the complex spike itself.

A potentially unsatisfactory aspect of our simulations was the time taken to learn. Of the order or 100 000 iterations were required to optimise the 400 independent variables of the cerebellar output. Stochastic gradient descent is inherently slow, since just one or a few of those variables can be perturbed in one movement realisation, and the weight changes are furthermore individually small. Before considering possible acceleration methods, we note that some motor behaviours are repeated huge numbers of times. An obvious example is locomotion. Thus, public health campaigns in vogue in the USA at the time of writing aim for people to take 10 000 steps per day. So, clearly, a target of a few hundred thousand steps could be achieved in a matter of days or weeks.

Part of the slowness of learning results from the conflicting pressures on the plastic weight changes. Large changes allow rapid learning, but could prevent accurate optimisation. An obvious extension of our implementation that would resolve this conflict would be to allow large plastic changes far from the optimum but to reduce them as the optimum is approached. The information required to do this is available as the net drive (error excitation – cancellation inhibition) to the olivary neurones at the time of emission of an error complex spike. If the drive is strong, one can imagine a long burst of action potentials being emitted. There is in vitro (Mathy et al., 2009) and in vivo (Rasmussen et al., 2013; Yang and Lisberger, 2014) evidence that climbing fibre burst length can influence plasticity in Purkinje cells. It seems possible that the same could be true in the cerebellar nuclei (or alternative plastic site in the subtraction of the average error). However, the above mechanism for adapting learning rates would only work directly in the LTD direction, since olivary cells cannot signal the strength of a net inhibition when no error complex spike is emitted.

A mechanism that could regulate the speed of learning in both LTP and LTD directions would be to target perturbations to the time points where they would be most useful—those with the greatest errors. This might be achieved by increasing the probability (and possibly the strength) of the perturbation complex spikes shortly before strongly unbalanced (excitatory or inhibitory) inputs to the olive. This process offers a possible interpretation for various observations of complex spikes occurring before error evaluation: related to movements (Bauswein et al., 1983; Kitazawa et al., 1998) or triggered by conditioned stimuli (Rasmussen et al., 2014; Ohmae and Medina, 2015).

Movement-specific adaptations of the learning rates could provide an explanation for the phenomenon of ‘savings’, according to which relearning a task after extinction occurs at a faster rate than the initial learning. The adaptations could plausibly be maintained during extinction and therefore persist until the relearning phase. These adaptations could appear to represent memories of previous errors (Herzfeld et al., 2014).

Finally, the output convergence solution we proposed above for the bicycle problem could also reflect a parallelisation strategy enabling the computations involved in stochastic gradient descent to be scaled from the small circuit we have simulated to the whole cerebellum. As mentioned above, this would probably require one of the above schemes for adjusting learning rates in a way that would allow plasticity in regions with ‘useful’ error information to dominate changes in those without.

### 9.6 Insight into learning in other brain regions

We believe that our proposed implementation of stochastic gradient descent offers possible insight into learning processes in other brain regions.

To date, the most compelling evidence for a stochastic gradient descent mechanism has been provided in the context of the acquisition of birdsong. A specific nucleus, the ‘LMAN’ has been shown to be responsible for song variability during learning and also to be required for learning (Doya and Sejnowski, 1988; Olveczky et al., 2005). Its established role is therefore analogous to our perturbation complex spike. Our suggestion that the same input (the climbing fibre) signals both perturbation and error change may also apply in the birdsong context, where it would imply that LMAN also assumes the role of determining the sign of plasticity at the connections it perturbed. However, such an idea has for now not been examined and there is as yet a poor understanding of how the trial song is evaluated and of the mechanism for transmitting that information to the adaptive site; indeed the adaptive site itself has not been identified unequivocally.

We see a stronger potential analogy with our mechanism of stochastic gradient descent in the learning of reward-maximising action sequences by the basal ganglia. Under the stimulus of cortical inputs, ensembles of striatal medial spiny neurones become active, with the resulting activity partition determining the actions selected by disinhibition of central pattern generators through inhibition of the globus pallidus (Grillner et al., 2005). It is thought that the system learns to favour the actions that maximise the (discounted) reward, which is signalled by activity bursts in dopaminergic midbrain neurones and phasic release of dopamine, notably in the striatum itself (Schultz, 1986). This has been argued (Schultz et al., 1997) to reflect *reinforcement learning* or more specifically *temporal difference learning* (Sutton and Barto, 1998).

We note that temporal difference learning can be decomposed into two problems: linking actions to potential future rewards and a gradient ascent to maximise reward. In respect of the gradient ascent, we note that dopamine has a second, very well known action in the striatum: it is necessary for the initiation of voluntary movements, since reduction of dopaminergic input to the striatum is the cause of Parkinson’s disease, in which volitional movement is severely impaired. The key point is to combine the two roles of the dopaminergic system in the initiation of movement and in signalling reward. The initiation of movement by dopamine, which could contribute to probabilistic action selection, would be considered analogous to our perturbation complex spike and could create an eligibility trace. A subsequent reward signal would result in plasticity of eligible synapses reinforcing the selection of that action, with possible sites of plasticity including the cortico-striatal synapses that were successful in exciting an ensemble of striatal neurones^8^. This would constitute a mechanism of gradient ascent analogous to that we have proposed for gradient descent in the cerebellar system.

Of particular interest is whether correct optimisation also involves a mechanism for subtracting the average error in order to extract gradient information. Such subtraction would be entirely consistent with the reports of midbrain dopaminergic neurones responding more weakly to expected rewards and responding with sub-baseline firing to omission of a predicted reward. These phenomena moreover appear to involve an adaptive inhibitory mechanism (Eshel et al., 2015). These observations could be interpreted as a subtraction of the average reward by a process analogous to that we propose for the extraction of the change of error 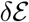.

### 9.7 Conclusion

We have proposed a complete and plausible mechanism of stochastic gradient descent in the cerebellar system, in which the climbing fibre perturbs movements, creates an eligibility trace, signals error changes and guides plasticity at the sites of perturbation. We verify predicted plasticity rules that contradict the current consensus and highlight the importance of studying plasticity under physiological conditions. The gradient descent requires extraction of the change of error and we propose an adaptive inhibitory mechanism for doing this via cancellation of the average error. Our implementation of stochastic gradient descent suggests the operation of an analogous mechanism (of gradient ascent) in the basal ganglia initiated and rewarded by dopaminergic signalling.

## Acknowledgements

We are grateful to the following for discussion and/or comments on this manuscript: David Attwell, Mariano Casado, Paul Dean, Anne Feltz, Richard Hawkes, Clément Léna, Steven Lisberger, Tom Ruigrok, John Simpson, Brandon Stell, Stéphane Supplisson and German Szapiro.

This work was funded by ANR-08-SYSC-005 (BB), the Labex MEMOLIFE (BB), the FRM DEQ20160334927 (BB) and NSF IIS-1430296 (JA, NB). GB was funded by Région Île-de-France, FRM, and Labex MEMOLIFE. JR received funding from the Deutsche Forschungsgemeinschaft (DFG, RA-2571/1-1). This work was supported by the program ‘Investissements d’Avenir’ from the French Government, implemented by Agence Nationale de la Recherche (ANR), references: ANR-10-LABX-54 MEMOLIFE and ANR-11-IDEX-0001-02 PSL* Research University.

## Author contributions

Concept BB; experimental design BB and GB; all experiments shown GB; all analysis shown BB; network simulation BB, JR; theoretical analysis VH, JA and NB, with help from J-PN and CC; pilot experiments GB and CB; suggestion of tracking plasticity in the cerebellar nuclei CB; analysis framework AB; manuscript BB, VH, GB, JA and NB.

## 10 Changes

Some minor textual changes are not listed.

### 10.1 Version 21-Nov-2016

1. The concentration of K-gluconate in the pipette solution was corrected to 128 mM from 148 mM, a typographical error.
2. In Fig. 3A and the associated text, ‘CS rate’ was introduced to represent climbing fibre acticity signalling a cell-and-movement-specific error signal and to distinguish that from the global error 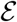. The legend and corresponding text have been modified appropriately.
3. The largest *n* in the analysis of paired-pulse ratios was corrected to 18 from 15.
4. Added reference to Suvrathan et al. (2016).
5. It is now specified that female mice were used and we have added statement about animal experimentation authorisation.

### 10.2 Version 29-Jul-2017

1. The description of the connectivity between mossy fibres and nucleo-olivary neurones was clarified.
2. Extended analysis of storage capacity and comparison with theoretical maximum, including a new Fig. 13, in collaboration with new author Johnatan Aljadeff.
3. A small change—adding a constant inhibition in nucleo-olivary neurones—prevented storage capacity for the error estimate from limiting learning.
4. References demonstrating that climbing fibre stimulation elicits movements have been added.
5. Added reference to Howe and Dombeck (2016)

### 10.3 Version 02-Oct-2017

1. Eqn. (37) was corrected.

### 10.4 Version 25-July-2018

1. Antonin Blot and Jonas Ranft were added as authors.
2. EGTA was incorrectly omitted from the recipe describing the pipette solution; the concentration used is now indicated.
3. Correction of a bug in the analysis has caused small changes to the plasticity time courses in Fig. 7. This was without effect on the ratios used in the statistical analysis.

## Footnotes

* The first version of this preprint was posted on 16-May-2016. Significant changes are listed in section §10.

1 The objective of decorrelation may appear at odds with the extensive literature reporting anti-correlation of Purkinje simple and complex spike modulations (e.g. Barmack and Yakhnitsa, 2003). This contradiction may only be apparent, because the decorrelation should apply to random variations of the discharge of parallel and climbing fibres about their means. Furthermore, most reports of anti-correlation were not obtained under conditions of execution of a learnt movement; more often they involve unusual stimuli to which the animal is naive, and the preparations were often anæsthetised or decerebrate.

2 This saccade data was chosen to illustrate the general problems facing cerebellar learning, but for this particular behaviour it should be noted that results that appear remarkably divergent have been reported in learning studies (reviewed in Dash and Thier, 2014; Soetedjo et al., 2008).

3 In reality, higher-level representations appear to be employed (Ebner et al., 2011).

4 For simplicity, we here consider only constant perturbation and learning steps. Therefore, at best the rate and the error fluctuates around the target rates and zero error, in a bounded domain of size determined by the magnitude of the constant perturbation and learning steps. We say that the algorithm converges when this situation is reached.

5 In the triangular domain below the line *C*_−_, the mean leftward-upward drift has an angle greater than the 3*π*/4 inclination of the *C*_−_ line for 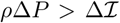 (i.e. *β* < *ρ*). In this case, the mean drift does not ensure that the 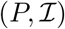 trajectory crosses the *C*_−_ line. However, if *P* becomes zero before crossing *C*_−_, the positivity constraint on the rate, imposes that subsequent updates are strictly upward, until 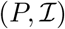 reaches *C*_−_.

6 More accurately, we neglect the number of crossings of the *C*_+_ line which is exponentially smaller in *A*/sup 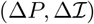 than the number of crossings of the *D*_+_ line.

7 For a fraction of the trajectories, the rate of convergence is about half of the prediction. This arises because these simulations constrained synaptic weights to be non-negative, which creates a significant fraction of synapses with negligible weights (Brunel et al., 2004) and produces a smaller effective step Δ*P* than estimated without taking this positivity constraint into account.

8 Rui Costa made a similar suggestion at the 5th Colloquium of the Institut du Fer à Moulin, Paris, 2014.

